# Unveiling the Core Functional Networks of Cognition: An Ontology-Guided Machine Learning Approach

**DOI:** 10.1101/2024.04.02.587855

**Authors:** Guowei Wu, Zaixu Cui, Xiuyi Wang, Yi Du

## Abstract

Deciphering the functional architecture that underpins diverse cognitive functions is fundamental quest in neuroscience. In this study, we employed an innovative machine learning framework that integrated cognitive ontology with functional connectivity analysis to identify brain networks essential for cognition. We identified a core assembly of functional connectomes, primarily located within the association cortex, which showed superior predictive performance compared to two conventional methods widely employed in previous research across various cognitive domains. Our approach achieved a mean prediction accuracy of 0.13 across 16 cognitive tasks, including working memory, reading comprehension, and sustained attention, outperforming the traditional methods’ accuracy of 0.08. In contrast, our method showed limited predictive power for sensory, motor, and emotional functions, with a mean prediction accuracy of 0.03 across 9 relevant tasks, slightly lower than the traditional methods’ accuracy of 0.04. These cognitive connectomes were further characterized by distinctive patterns of resting-state functional connectivity, structural connectivity via white matter tracts, and gene expression, highlighting their neurogenetic underpinnings. Our findings reveal a domain-general functional network fingerprint that pivotal to cognition, offering a novel computational approach to explore the neural foundations of cognitive abilities.

## Introduction

Unraveling the complexity of the human mind necessitates an in-depth examination of its cognitive functions such as attention, memory, executive functions, language, social cognition, all intertwined and foundational to our understanding of neuroscience [1–3]. The advent of cognitive ontology has paved the way for a structured exploration of these mental capacities, mapping their intricate interrelations with a data-driven approach [4–8]. Despite advancements in the construction of cognitive ontology, revealing its neural underpinnings remains a significant challenge.

Recent studies have leveraged functional connectivity (FC)—the correlation between the BOLD time series of different brain regions—to predict cognitive abilities with notable success, offering insights into the brain’s organizational principles [9–12]. However, inconsistencies in methodology across studies have obscured a unified neural basis for cognitive ontology entity. For instance, Dubois et al. [10] utilized exploratory factor analysis (EFA) to establish a classical cognitive ontology entity, namely the g-factor, and demonstrated its predictability through distributed FC networks. In contrast, Kong et al. [11] applied factor analysis to develop a different cognitive ontology entity, the cognitive factor, without examining its predictability through FC networks. This disparity highlights the need for a coherent understanding of FC networks’ role in cognition prediction.

Emerging research have underscored specific brain networks’ pivotal role in supporting cognition. Bolt et al. [13] identified a general factor indicative of a “focused awareness” process or “attentional episode” through exploratory bifactor analysis, symbolizing a domain-general psychological process across cognition, perception, action, and emotion domains, manifest in core regions of the task-positive network. In parellel, Kristanto et al. [14] investigated the alignment between brain-derived (neurometric) and cognitive performance-derived (psychometric) ontologies, revealing that regions associated with cognitive abilities form an extensively interconnected structural network. These findings underscore the significance of selected brain regions for cognitive functions, laying the groundwork for our hypothesis that certain FC networks are crucial in supporting cognition.

Further bolstering this hypothesis, studies have demonstrated the cognitive factor’s predictive power across multiple cognitive domains and its correlation with individual differences in ideological preferences [1,3,5,6]. He et al.’s [15] meta-matching framework, applying predictive models from large-scale datasets (the UK Biobank, N = 36,848) to new, smaller-scale (the Human Connectome Project, HCP, N = 1,019) non-brain-imaging phenotypes, has further exemplified the predictive capacity of FC data. Motived by these findings, we aimed to explore FC networks’ capacity to encapsulate the cognitive factor and predict diverse cognitive functions beyond initial study groups.

Given the cognitive factor’s stronger link to cognitive functions over sensory, motor, or emotional functions [7,16] we posited that distinct FC networks would preferentially represent cognitive abilities like executive function and language, but not those less cognitively related, such as motor function and emotion. Furthermore, we hypothesize that the FC networks crucial for representing the cognitive factor may primarily consist of connections within the association cortex. The association cortex, encompassing regions such as the prefrontal cortex, parietal cortex, and temporal cortex, has been implicated in higher-order cognitive processes, including attention, working memory, decision-making, and language. These regions exhibit high levels of functional connectivity, forming a densely interconnected network that supports the integration and processing of information [17–20]. Moreover, the cognitive factor, as a latent variable derived from diverse cognitive tasks, likely reflects a domain-general cognitive ability that underlies performance across multiple domains [1,3,5,6,15]. As such, the FC networks that effectively represent the cognitive factor should be able to predict individual differences in various cognitive functions, including attention, working memory, executive control, and language processing. This hypothesis aligns with recent findings demonstrating the predictive power of FC data across different cognitive domains and datasets [10,11,16].

Utilizing behavioral and resting-state fMRI data from the HCP Young Adult (HCP-YA) [21] and HCP Development Study (HCP-D) [22], we sought to validate if specific functional networks could consistently predict the cognitive factor constructed by different methods, and subsequently, cognitive behaviors. This approach not only tested the predictive stability of these networks but also delved into the biological basis of FC networks associated with the cognitive factor, examining FC strength variability, white matter connectivity, and gene association patterns. Our comprehensive approach endeavors to elucidate the complex interplay between functional and structural connectivity, gene expression, and cognition, ultimately enriching our understanding of the neural foundations of human intelligence and behavior.

## Method

### (a) Participant information

We used two datasets: the WU-Minn HCP-YA S1200 release [21] and HCP-D [22]. The HCP-YA dataset was used to test the main hypothesis: whether a unique group of network FC edges underlying the cognitive factor can effectively predict various cognitive related behaviors rather than cognitive less-related behaviors. The HCP-D dataset was employed to evaluate the capacity of this unique network configuration to predict cognitive behaviors across datasets with comparable efficacy. Please visit HCP project’s website (https://www.humanconnectome.org/) for detailed information about data acquisition protocols. This project was approved by the Washington University local ethics committee.

We utilized the publicly accessible data from the HCP-YA S1200 release, encompassing multiple magnetic resonance imaging (MRI) modalities such as resting-state functional MRI, diffusion MRI, and structural MRI. The imaging data were obtained using a Siemens 3T Skyra scanner with a multiband sequence. Each participant underwent 2 separate fMRI sessions on different days, with each session comprising 2 phase-encoding runs (left-right and right-left, LR/RL). T1-weighted structural data involved a single 0.7 mm isotropic scan per participant. The dMRI data encompassed six 1.25 mm isotropic runs for each participant. Further information about the dataset and MRI acquisition parameters can be found in the prior study by [21]

In addition to imaging data, the HCP database furnishes demographic data and performance metrics for numerous cognitive tasks. From the available 1,206 datasets within the HCP database at the time of the study, we excluded those with incomplete information, insufficient fMRI images, motion unqualified data, and left-handed individuals. Consequently, a final sample of 601 individuals (329 females; aged between 22 and 36 years), was included.

This study incorporated a total of 633 subjects (294 males, aged 8-21 years) from the Lifespan 2.0 Release of the HCP-D. All imaging data were collected utilizing a multiband EPI sequence on a 3T Siemens Prisma scanner. Each participant took part in 2 separate rs-fMRI sessions on different days.. The study employed a total of 4 runs, accounting for 26 minutes of rs-fMRI. T1-weighted structural data involved a single 0.8 mm isotropic scan per participant. For further information about the dataset and MRI acquisition parameters, please refer to [22].

Participants were excluded including those with incomplete information, insufficient fMRI images, motion unqualified data, and left-handed individuals. The final sample thus comprised 374 individuals (197 females; age: 8 - 22).

### (b) The pre-processing of fMRI data and functional connectivity estimation

In this study, we employed minimally preprocessed T1w and fMRI images from both HCP-D and HCP-YA. The T1w images were initially corrected for intensity non-uniformity and subsequently used as T1w-reference throughout the workflow. Following skull-stripping and brain surface reconstruction, the volume-based brain-extracted T1w images were segmented into cerebrospinal fluid (CSF), white matter (WM), and gray matter (GM), and spatially normalized to MNI space. The rs-fMRI images underwent preprocessing steps such as motion correction, distortion correction, co-registration to the T1w-reference image, and normalization to MNI space [23]. The BOLD timeseries were resampled to the fsaverage space, generating grayordinates files containing 91k samples.

Subsequently, the preprocessed fMRI images underwent further post-processing using the extensible Connectivity Pipelines (XCP-D) [24]. Prior to nuisance regression, volumes with framewise-displacement (FD) exceeding 0.3 were flagged as outliers and excluded. A total of 36 nuisance regressors were regressed out from the BOLD data, including motion parameters, global signal, mean white matter, and mean CSF signal with their temporal derivatives, as well as the quadratic expansion of motion parameters and tissue signals with their derivatives. The residual timeseries were then band-pass filtered (0.01-0.08 Hz) and spatially smoothed (FWHM = 6 mm) [25,26].

To mitigate the potential impact of head motion, subjects were further excluded based on two criteria [27,28]. First, we excluded fMRI data with more than 25% of FDs exceeding 0.2 mm. Second, we excluded fMRI data if the mean FD surpassed the 1.5*interquartile range (IQR) in the adverse direction when estimating the distribution of mean FD across the same run for each dataset. We retained subjects with all fMRI runs in HCP-YA (4 rs-fMRI runs) and HCP-D (4 rs-fMRI runs). A total of 601 subjects (329 females, age 22-36) in HCP-YA and 374 subjects (197 males, age 8-22) in HCP-D were utilized in the subsequent analyses.

We employed the Schaefer parcellation [29] to extract regional Blood Oxygen Level Dependent (BOLD) timeseries data. This parcellation divides the whole brain into 400 parcels and has been extensively utilized [30–32]. Previous studies showed commendable performance when using this parcellation in the context of FC network analysis to predict cognitive functions[11,33,34]. We evaluated FC by calculating the Pearson correlation coefficient between the timeseries of parcels. As a result, we obtained a symmetric 400 * 400 functional connectivity matrix for each participant. This matrix was transformed using the Fisher transformation, and the upper triangle was vectorized, generating 79800 connectivity features.

### (c) Cognitive factor computation

To derive a psychometric ontological entity, we conducted a confirmatory factor analysis (CFA) model using the behavioral data from 13 tasks of the HCP-YA dataset. CFA can robustly construct ontological entities [3,10,14]. We selected 13 behavioral metrics adopted from the task-evoked fMRI measurement (in-scanner test), NIH Toolbox, and Penny Cognitive Battery based on previous studies [3,10,14]. These behavioral metrics cover 5 distinct cognitive domains, including: (1) executive functions related to processing speed: the NIH unadjusted scale score for Dimensional Change Card Sort, Flanker test and Pattern completion Processing Speed, (2) executive functions associated with working memory: Reaction time of 2-back condition in the working memory test, the NIH unadjusted scale score for List Sorting task and Picture Sequence Memory, (3) fluid intelligence: Number of Correct Responses (PMAT24 A CR), Total Skipped Items (PMAT24 A SI), and Median Reaction Time for Correct Responses (PMAT24 A RTCR) in Penn Progressive Matrices test, (4) relation ability: Reaction time of Relation task of match condition, Reaction time of Relation task of relevance condition and (5) language ability: the NIH unadjusted scale score for Picture Vocabulary test, and Reading Recognition test. A summary of these tasks is provided in Table 1.

**Table 1.**
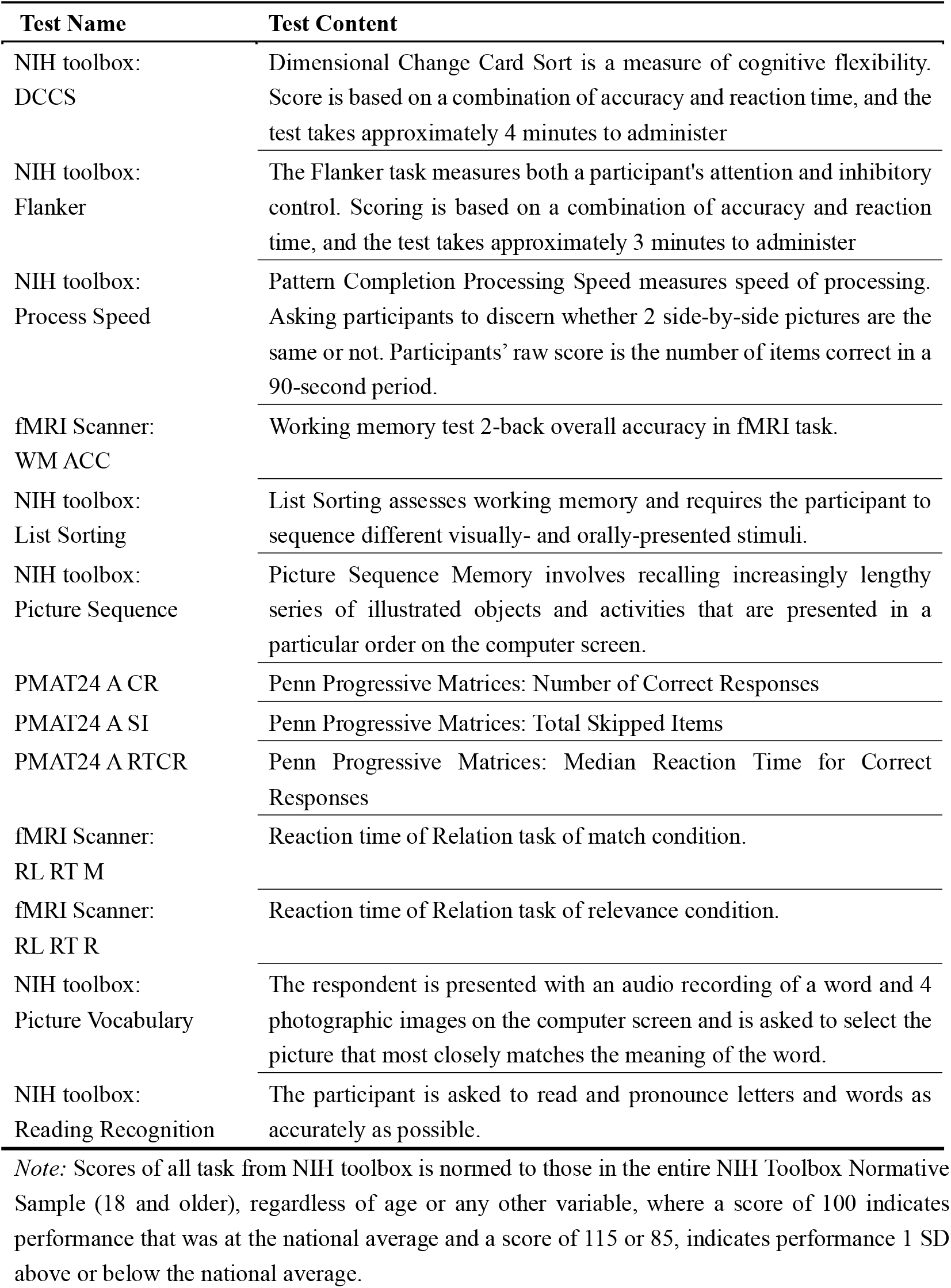
13 behaviors used in CFA model.

**Table 2.**
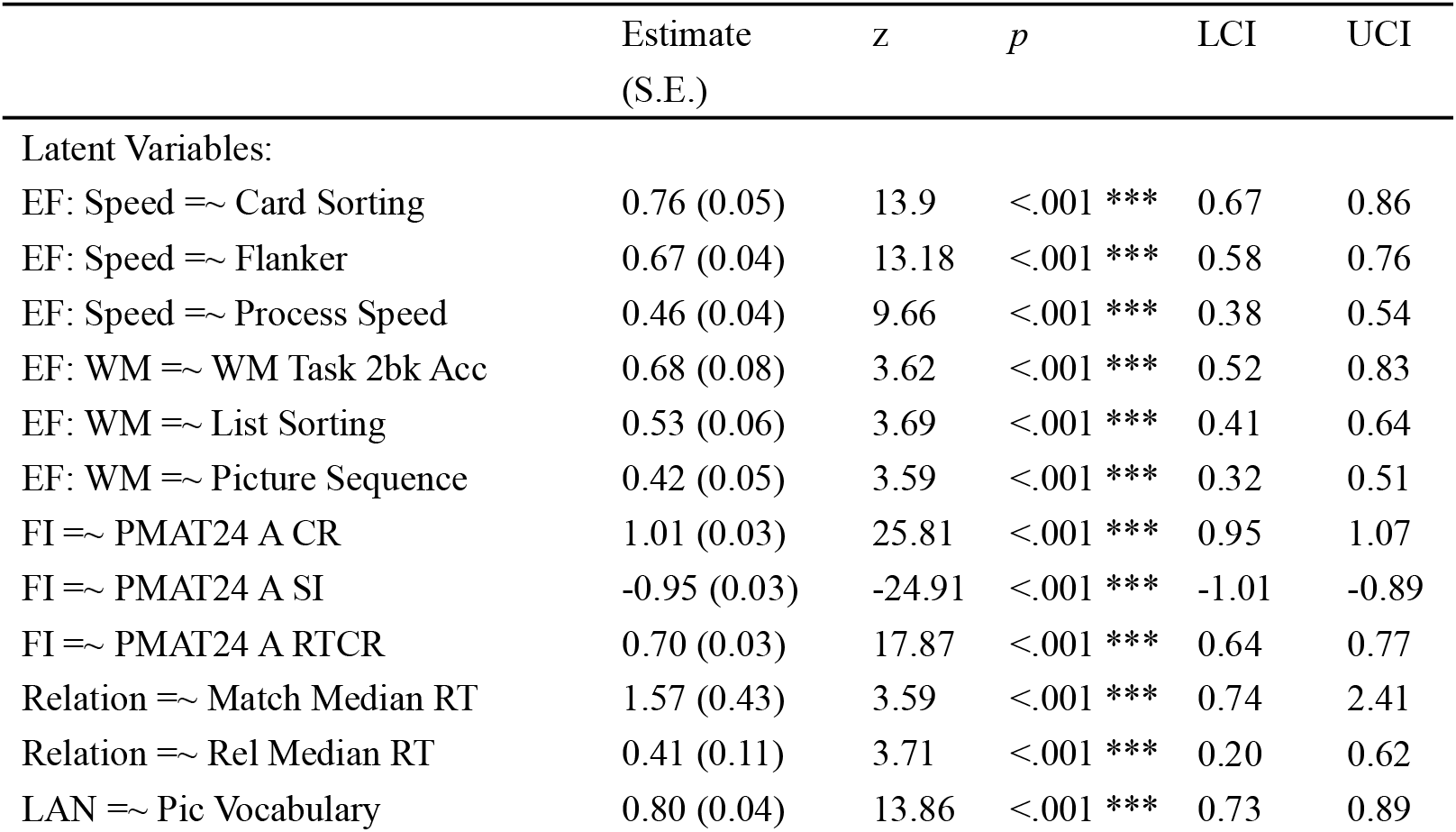
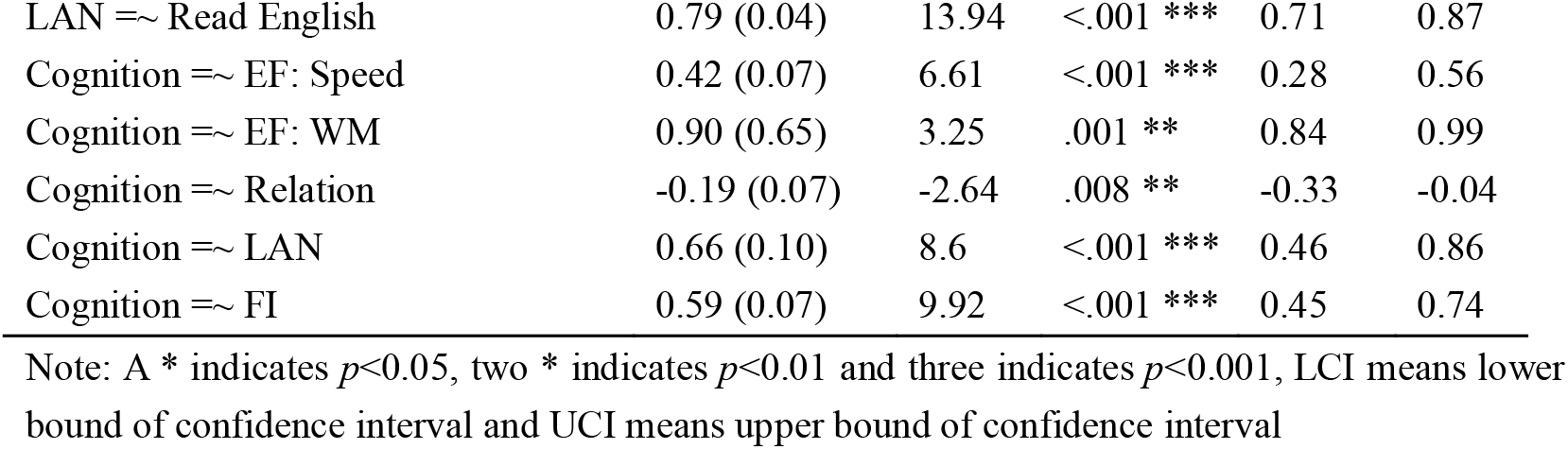
Fitting results of second order CFA model.

|  | Estimate<br>(S.E.) | z | p | LCI | UCI |
| --- | --- | --- | --- | --- | --- |
| Latent Variables: |  |  |  |  |  |
| EF: Speed =~ Card Sorting | 0.76 (0.05) | 13.9 | <.001 *** | 0.67 | 0.86 |
| EF: Speed =~ Flanker | 0.67 (0.04) | 13.18 | <.001 *** | 0.58 | 0.76 |
| EF: Speed =~ Process Speed | 0.46 (0.04) | 9.66 | <.001 *** | 0.38 | 0.54 |
| EF: WM =~ WM Task 2bk Acc | 0.68 (0.08) | 3.62 | <.001 *** | 0.52 | 0.83 |
| EF: WM =~ List Sorting | 0.53 (0.06) | 3.69 | <.001 *** | 0.41 | 0.64 |
| EF: WM =~ Picture Sequence | 0.42 (0.05) | 3.59 | <.001 *** | 0.32 | 0.51 |
| FI =~ PMAT24 A CR | 1.01 (0.03) | 25.81 | <.001 *** | 0.95 | 1.07 |
| FI =~ PMAT24 A SI | -0.95 (0.03) | -24.91 | <.001 *** | -1.01 | -0.89 |
| FI =~ PMAT24 A RTCR | 0.70 (0.03) | 17.87 | <.001 *** | 0.64 | 0.77 |
| Relation =~ Match Median RT | 1.57 (0.43) | 3.59 | <.001 *** | 0.74 | 2.41 |
| Relation =~ Rel Median RT | 0.41 (0.11) | 3.71 | <.001 *** | 0.20 | 0.62 |
| LAN =~ Pic Vocabulary | 0.80 (0.04) | 13.86 | <.001 *** | 0.73 | 0.89 |
| LAN =~ Read English | 0.79 (0.04) | 13.94 | <.001 *** | 0.71 | 0.87 |
| Cognition =~ EF: Speed | 0.42 (0.07) | 6.61 | <.001 *** | 0.28 | 0.56 |
| Cognition =~ EF: WM | 0.90 (0.65) | 3.25 | .001 ** | 0.84 | 0.99 |
| Cognition =~ Relation | -0.19 (0.07) | -2.64 | .008 ** | -0.33 | -0.04 |
| Cognition =~ LAN | 0.66 (0.10) | 8.6 | <.001 *** | 0.46 | 0.86 |
| Cognition =~ FI | 0.59 (0.07) | 9.92 | <.001 *** | 0.45 | 0.74 |

Note: A \* indicates $p < 0.05$ , two \* indicates $p < 0.01$ and three indicates $p < 0.001$ , LCI means lower bound of confidence interval and UCI means upper bound of confidence interval

There are several methods to derive a cognitive ontological factor from scores on a set of cognitive tasks, and a consensus comprehensive framework is yet to be established to define cognitive ontology entity [7,8,16]. However, previous research indicated that cognitive domains, such as executive functions, fluid intelligence, and language abilities, are not independent but rather exhibit a hierarchical cognitive organization [3]. Accordingly, we proposed a second-order CFA model to construct a domain-general cognitive ontological entity. To achieve this, we utilized the *lavaan* package (https://lavaan.ugent.be) within the R Software (https://www.r-project.org/).

We hypothesized that all 13 task scores can be organized into the five first-order factors: executive function related to process speed, executive function related to working memory, relation ability, fluid intelligence, and language ability [3,10,14]. Additionally, we proposed a second order latent variable called ‘cognitive factor’ to capture the interrelationships among these five first-order factors according to EFA model result from previous study [3]. We fitted this model to the data of HCP-YA which included a total of 601 subjects. To facilitate the model identification, all behavior scores were standardized using z-scores, allowing for free estimation of all factor loadings. The CFA model was tested using Maximum Likelihood (ML) estimator, and the model fit was assessed using the chi-square goodness-of-fit index (χ2), the comparative Fit Index (CFI), the Root-Mean-Squared Error of Approximation (RMSEA), and the Standardized Root-Mean-Squared residual (SRMR). We evaluated the model fit based on established thresholds, considering CFI equal to or greater than 0.95 and both RMSEA and SRMR values less than 0.08, as recommended by [35] After model fitting, we calculated the individual-level cognitive factor score with a regression method using *lavPredict* function [36].

### (d) Prediction of the cognitive factor score using FC

To identify the FC edges that have a significant influence on predicting cognitive factor score, we employed ridge regression to examine the relationship between all FC edges and cognitive factor score obtained from the CFA model [11,17,20,37]. We set the second-order cognitive factor score as the dependent variable, enabling us to calculate the regression coefficient (β value) for each edge corresponding to the cognitive factor score. The L2-norm regularization of ridge regression archived a balance between penalties for training error and model complexity. The magnitude of the β coefficient for each edge indicates its significance in predicting the cognitive factor score. We sorted all the absolute weights of the β coefficient and selected the top 10% ranked edges. In this way, we identified a subset of FC edges vital for predicting the cognitive factor score. To train the ridge regression model, we employed a nested 2-fold cross-validation strategy. For each iteration, we divided the dataset into two parts for training and validation [17]. Specifically, we measured prediction accuracy using the Pearson correlation and Mean Squared Error (MSE) between actual and predicted values. For each iteration, roles of training and testing sets were switched, producing two Pearson R values and MSE values. We averaged these values across 100 iterations to determine the model’s overall performance. The term “prediction accuracy,” when not specifically defined, refers to the Pearson correlation coefficient ‘R’ between predicted and actual values.

To assess if FC edges significantly predict cognitive factor score, we established a null distribution using a permutation test. This involved randomly shuffling cognitive factor scores within the training dataset across participants 200 times, creating 200 samples of prediction accuracy under the assumption of no relationship between FC and cognitive ontology score. We then compared the 100 actual correlation coefficients (R values) from the above 100 iterations with this null distribution using the nonparametric Wilcoxon signed-rank test [12,20].

### (e) Behaviors used in cross task prediction paradigm

To assess whether the key FC edges used to predict cognitive factor score accurately represent cognitive functions, we chose 25 behavioral metrics from the HCP-YA dataset in a cross-task prediction paradigm. These target behaviors encompass a variety cognitive related behavior, such as working memory, fluid intelligence, language ability, spatial attention, and sustained attention. We also included several cognitive less-related behaviors, such as motor tasks, sophisticated emotional processing tasks, and measures assessing life satisfaction. These behaviors allow us to evaluate if the pivotal FC edges for predicting cognitive factor score can better forecast cognitive functions compared to motor or emotion functions.

The 25 selected behavioral measures span 10 domains, including:

1. one metric for Working Memory: Median Reaction Time of 2-back condition of Working Memory Task in fMRI scanner (WM Task 2bk Median RT);
2. three metrics for Language ability: Accuracy of Language Task in Story condition (Language Task Story Acc), Average Difficulty in Language Task in Story condition (Story Avg Difficulty) and Median Reaction Time of Language Task in Story condition (Language Task Story Median RT);
3. three metrics for Sustained Attention: Learning Rate in Short Penn Continuous Performance Test (SCPT LRNR), Sensitivity in Short Penn Continuous Performance Test (SCPT SPEC), Specificity in Short Penn Continuous Performance Test (SCPT SEN) and Median Response Time for True in Short Penn Continuous Performance Test (SCPT TPRT);
4. three metrics for Spatial Orientation: Correct Reaction Time in Penn Line Orientation Test (VSPLOT CRTE), Offset in Penn Line Orientation Test (VSPLOT OFF) and Total completion in Penn Line Orientation Test (VSPLOT TC);
5. one metric for Self-Regulation: Delay Discounting: Area Under the Curve for Discounting of $40,000 (DDisc AUC 40K);
6. one for Verbal Episodic Memory: Median Reaction Time for Correct Responses in Penn Word Memory Test (IWRD RTC);
7. one metric for Psychological Well-being: Subjective well-being and contentment with life (Life Satisfy);
8. four metrics for Social Relationship: Perceived interpersonal rejection (Rejection), Subjective feelings of loneliness and social isolation (Loneliness), Perceived helpful behaviors from others (Instru Supp) and Perceived stress level (PercStress);
9. three metrics for Emotion: Intensity of Anger emotions (Anger Affect), Intensity of Sadness emotions (Sadness) and Intensity of Fear emotions (Fear Affect);
10. four metrics for Motor: Endurance, Grip Strength (Strength), Walking Speed (Gait Speed Component) and Dexterity.

Detailed descriptions of each measure are available in Table 3. All task scores were standardized using z-scores before being included in the prediction model. To measure what the extent that cognitive factor related to these 25 target behaviors, we calculated the Spearman correlation between cognitive factor score and their raw scores, which we refer to as ‘cognitive loading’. Behaviors that showed a significant correlation with cognitive factor are considered cognitive-related behaviors, while others are categorized as cognitive less-related behaviors.

**Table 3.**
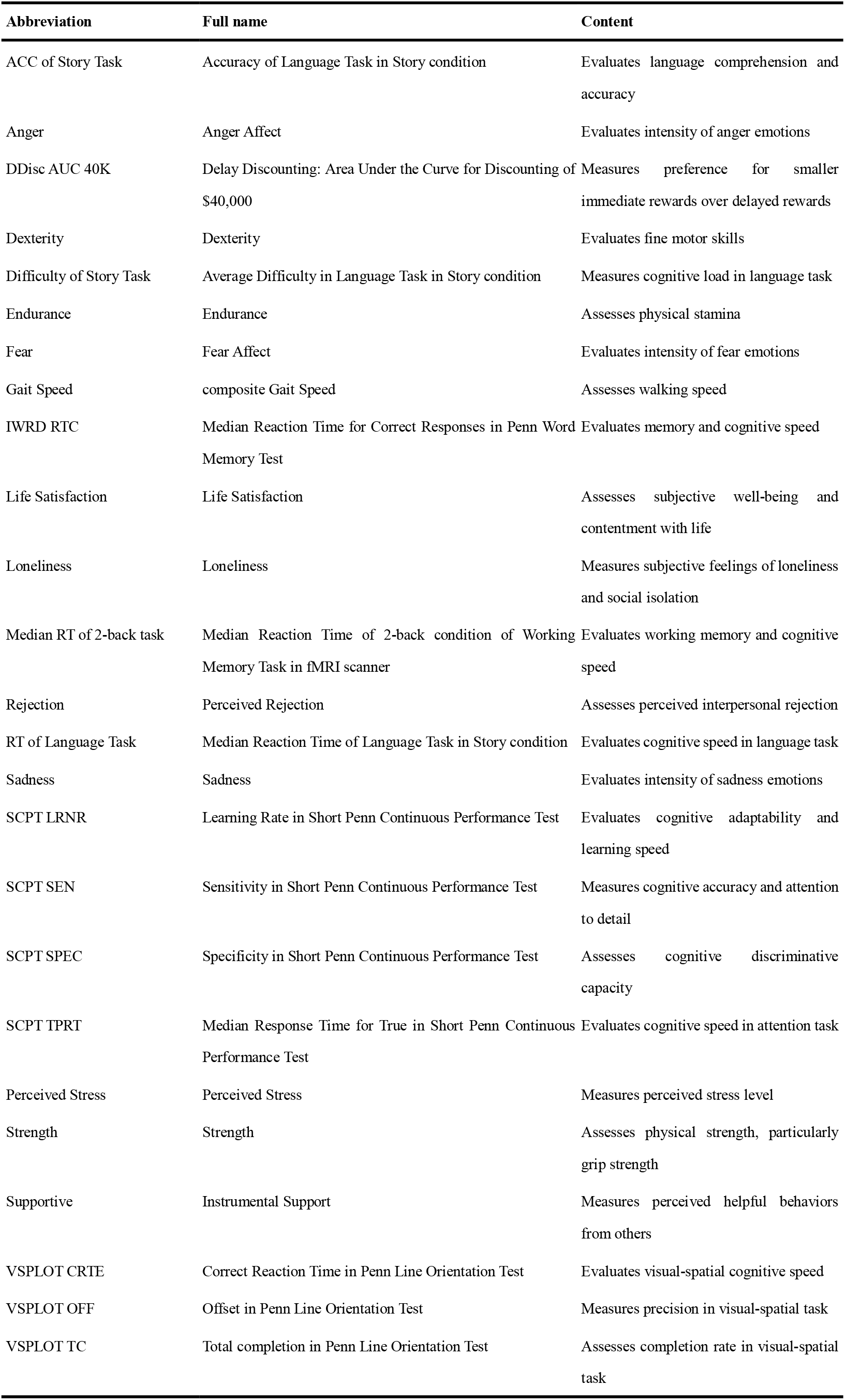
Detailed descriptions of 25 tasks from HCP-YA used in prediction analysis.

**Table 4.** The prediction accuracy of all behaviors for all three models.

| Domains | Behavior | Full FC (SD) | rCPM (SD) | COPM (SD) |
| --- | --- | --- | --- | --- |
| Executive function | WM Task 2bk Median RT | 0.084 (0.044) | 0.102 (0.033) | 0.137 (0.043) |
| Emotion | Anger Affect | 0.044 (0.044) | 0.049 (0.030) | 0.028 (0.037) |
| Emotion | Sadness | 0.038(0.043) | 0.064 (0.030) | 0.024 (0.038) |
| Emotion | Fear Affect | 0.031(0. 038) | 0.043 (0. 030) | 0.008 (0. 041) |
| Language | Language Task Story Avg<br>Difficulty | 0.189 (0.031) | 0.194 (0.029) | 0.346 (0.030) |
| Language | Language Task Story Acc | 0.048 (0.046) | 0.094 (0.037) | 0.136 (0.042) |
| Language | Language Task Story Median RT | 0.095 (0.037) | 0.060 (0.034) | 0.107 (0.037) |
| Psychological<br>Well-being | Life Satisfy | 0.072 (0.039) | 0.060 (0.032) | 0.113 (0.034) |
| Motor | Endurance | 0.107 (0.043) | 0.130 (0.034) | 0.147 (0.043) |
| Motor | Strength | 0.053 (0.041) | 0.130 (0.034) | 0.055 (0.036) |
| Motor | Gait Speed COPM | 0.046 (0.046) | 0.049 (0.034) | 0.036 (0.043) |
| Motor | Dexterity | -0.038 (0.043) | 0.009 (0.030) | 0.026 (0.041) |
| Sustained Attention | SCPT TPRT | -0.007 (0.038) | 0.071 (0.044) | 0.116 (0.040) |
| Sustained Attention | SCPT SPEC | -0.002 (0.038) | 0.019 (0.032) | 0.070 (0.038) |
| Sustained Attention | SCPT SEN | 0.020 (0.039) | 0.048 (0.028) | 0.050 (0.039) |
| Sustained Attention | SCPT LRNR | -0.005 (0.033) | -0.016 (0.025) | -0.017 (0.030) |
| Spatial Orientation | VSLOT OFF | 0.221 (0.035) | 0.122 (0.035) | 0.326 (0.032) |
| Spatial Orientation | VSLOT TC | 0.194 (0.039) | 0.195 (0.031) | 0.285 (0.033) |
| Spatial Orientation | VSLOT CRTE | 0.074 (0.033) | 0.090 (0.032) | 0.071 (0.041) |
| Self-Regulation | DDisc AUC 40K | 0.217 (0.039) | 0.172 (0.033) | 0.223 (0.036) |
| Social Relationship | Rejection | 0.023 (0.039) | -0.006 (0.030) | 0.019 (0.042) |
| Social Relationship | Loneliness | -0.005 (0.043) | 0.049 (0.033) | -0.009 (0.043) |
| Social Relationship | Supportive | 0.003 (0.041) | -0.005 (0.030) | -0.016 (0.040) |
| Social Relationship | Perceived Stress | 0.036 (0.042) | 0.019 (0.032) | 0.069 (0.040) |
| Verbal Episodic<br>Memory | IWRD RTC | -0.048(0.040) | -0.051 (0.031) | 0.002 (0.042) |

### (f) Framework of FC based behavioral prediction

We hypothesized that the edges that contributed significantly to predicting cognitive factor score could better represent cognitive functions compared to motor or emotion functions. To test this, we built three comparable models and then used these models to predict the behavior performance. Finally, we compared the prediction accuracy between these three models.

We built three models using identical cross-validation strategies, differing only in feature selection. The first named Cognitive Ontology based Prediction Model (COPM), selected FC features based on their importance in predicting cognitive factor score, using the top 10% of FC edges (7980) identified by their absolute β coefficients. This model, like the others, was applied to predict various behaviors, with each model’s prediction accuracy averaging over 100 iterations for stability. As baseline, we also included 2 additional models. The Full-FC (F-F) model, utilized all FC edges to predict behavioral score. To control feature number differences between COPM and F-F, our second baseline model, the Ridge-regression based Connectome Predictive Model (rCPM) also used 7980 edges. Unlike the COPM, which identifies edges based on the training dataset’s absolute β coefficients after 100 iterations of cognitive factor prediction modeling, the rCPM selects edges by ranking the absolute magnitude of each edge’s regression coefficients within the training set for specific behavioral prediction. For each model and each iteration, we used the split-half validation method, dividing the dataset into two parts for training and validation [17]. Specifically, we measured prediction accuracy using the Pearson correlation and MSE between actual and predicted values. For each iteration, roles of training and testing sets were switched, producing two Pearson R values and MSEs. We averaged these values across iterations to determine the model’s overall performance.

Finally, we assessed whether the COPM could better represent cognitive functions compared to motor or emotion functions, a distinction not observed in the F-F and rCPM. Specifically, we selected 25 behavioral tasks across 10 domains from the HCP-YA dataset. For each task, we first calculated the ‘cognitive loading’, which is the correlation between raw performance and cognitive factor score. Next, we calculated the average prediction accuracy for each model and calculated the Spearman correlations between the prediction accuracy and cognitive loading, labeling them as COPM-R, F-F Model-R, and rCPM-R, respectively. To determine if COPM more accurately predicts behaviors associated with cognitive factor score, we compared these correlations, using the *cocor* function in R [38,39]. A higher COPM-R value compared to both F-F Model-R and rCPM-R indicates that the COPM model offers more accurate predictions for behaviors closely associated with cognition. Furthermore, to demonstrate that COPM’s benefits are primarily for cognitive-related behaviors, we compared the mean prediction accuracy between tasks that are cognitive-related and those less so. This comparison was done using the non-parametric Wilcoxon signed-rank test, with all p-values adjusted using false discovery rate (FDR) correction.

For hyperparameter tuning, we employed a nested 20-fold cross-validation [20], specifically designed for ridge regression using MATLAB’s fitlinear.m function. In each inner training iteration, a grid search was performed on 100 logarithmically spaced α values ranging from 10^-5/N to 10^5/N, where N represents the number of subjects in the inner training fold. The optimization process combined Average Stochastic Gradient Descent (ASGD) with the limited-memory BFGS quasi-Newton algorithm (LBFGS), terminating after 1000 iterations or when the change in coefficients and intercept fell below 10^-4 per iteration. The optimal α value, yielding the minimum error across inner test folds, was used to calculate beta coefficients for the outer training set, while accounting for age, sex, and motion as covariates [11,20]. The resulting regression parameters were then applied to the testing set to control for covariate effects on behavioral variables.

### (g) Cross-dataset predictive efficiency of the COPM

We evaluated the COPM model’s ability to predict cognitive functions using the HCP-D dataset. This dataset includes similar behavioral tasks as HCP-YA, except that it focuses on children aged between 8 and 22. We hypothesized that if the COPM outperforms the F-F model and rCPM in predicting cognitive functions in the HCP-D dataset, while showing similar performance in motor and emotion functions, it would suggest a stable set of FC edges representing human cognition. For this test, we selected eight behavioral measures covering a broad range of domains. From our HCP-YA dataset analysis, 6 of these tasks assess cognitive related functions, such as executive control and language skills, while 2 measure cognitive less-related functions, including motor ability and emotional processing. The tasks include (1) 4 tasks for measuring executive function: Dimensional Card Sorting, Flanker test and List Sorting Working Memory and Picture Sequence Memory; (2) 2 tasks for measuring language ability, including Oral Reading Recognition test and Picture Vocabulary; (3) 1 task for motor: Grip Strength and (4) 1 task for emotion: Fear Affect, with detailed descriptions in S2 Table. We applied the COPM, F-F model, and rCPM to predict each behavior using the same procedures as before. We then used the non-parametric Wilcoxon signed-rank test to compare the prediction accuracy across the 2 types of behaviors and the 3 models. The COPM employed the top 10% of FC edges identified as significant in predicting cognitive factor score within the HCP-YA dataset. We did not reassess cognitive factor within the HCP-D dataset, thus directly assessing the cross-dataset predictive stability of these influential edges.

### (h) Exploring the FC variability, white matter structural basis, and genetic basis of FC edges in COPM

To explore the biological underpinnings of the 7980 FC edges used in the COPM, we analyzed their FC variability, white matter structural connectivity, and genetic expression similarity patterns, following the methodology outlined by [40]. To ascertain the significance of each edge in predicting cognitive factor, we calculated their linear contribution weights, called the ‘contribution weight matrix’, based on the cognitive factor prediction model. We obtained four 400x400 matrices, corresponding to contribution weight, FC strength variability, white matter structural connectivity, and genetic expression similarity, respectively.

The 400 parcels in the Schaefer 400 atlas can be classified into 17 functional networks [29,41]. We calculated intra-and inter-network connectivity averages within the Yeo 17 network [29,41] for each of the 400x400 matrices, yielding four distinct 17x17 matrices. Since we were only interested in the edges that make great contributions when predicting cognitive factor, we only considered these edges and calculated the intra-and inter-network average connectivity. After that, we computed the Spearman correlation between the contribution weight matrix and the other three matrices. To evaluate the statistical significance of these correlations, we generated a null distribution using permutation by randomly selecting 7980 edges 10000 times from the 400x400 matrix. We then computed intra-and inter-network connectivity averages within the Yeo 17 network, resulting in a new 17x17 matrix for each index. We subsequently correlated this matrix with the 17x17 contribution weight matrix. The *p*-value was determined using the proportion of real correlations that exceeded the permuted correlations, with a threshold for significance set at *p* < 0.05/3.

### (i) Validation analysis across varied edge selection percentiles and parcellation scheme

To assess the stability of our findings, we analyzed the most influential edges in predicting cognitive score using varying thresholds: 20%, 30%. For each threshold, we identified the edges with the greatest contribution and replicated the initial procedure. This process included predicting the performance of specific tasks at each threshold and then evaluating the correlation between cognitive loading and the prediction accuracy of both the COPM and rCPM models. Finally, we compared these correlation results across the 2 models for each threshold.

We further assessed our results’ robustness with Glasser’s atlas [42], constructing 360x360 individual functional connectomes to predict cognitive factor using COPM and F-F models. The F-F model alone served as the baseline for comparison, given its similar prediction accuracy to the rCPM model with the Schaefer atlas [29], and the time-consuming nature of replicating this analysis with rCPM.

## Code availability

All code used to perform the analyses in this study can be found at https://github.com/guoweiwuorgin/FC_COPM_framework.

## Data availability

The HCP-YA and HCP-D datasets, including T1-weight MRI, functional MRI and diffusion-weighted MRI are available at https://db.humanconnectome.org/. Gene expression data can be downloaded from the AHBA (http://human.brain-map.org) and processed using the abagen toolbox (https://github.com/rmarkello/abagen).

## Results

We first examined whether specific FC edges between networks could reliably represent the cognitive factor. This factor was constructed using CFA based on multiple behavior tasks from the HCP-YA dataset. We then calculated parcel-based FC matrices for each participant using resting-state fMRI data from the HCP-YA dataset, employing the Schaefer atlas which divides the cerebral cortex into 400 parcels [29]. Subsequently, we built a ridge regression model to identify FC edges crucial for predicting the cognitive factor. Next, we assessed whether a model incorporating these edges, the COPM outperformed two baseline models in predicting cognitive related behaviors whose performance showed significant correlations with the cognitive factor score, but not in predicting those cognitive less-related behaviors which showed no significant correlation with the cognitive factor score. The first baseline model, the F-F model, incorporated all edges, while the second used a connectome predictive model (CPM) method (termed as rCPM) to select the same number of edges as COPM but relevant to target behavior, not cognitive factor. We further validated the predictive reliability of these edges for cognitive functions using the HCP-D dataset, which includes 8 tasks identical to those in the HCP-YA. To probe the biological underpinnings of these key edges, we analyzed their relationships with FC variability, white matter structure, and gene expression similarity. Moreover, to ensure the robustness of our findings, we replicated the main results using varying thresholds for edge selection within the COPM, alongside using different brain parcellations to delineate functional areas. The whole analysis framework outlined in Figure 1.

**Figure 1.**
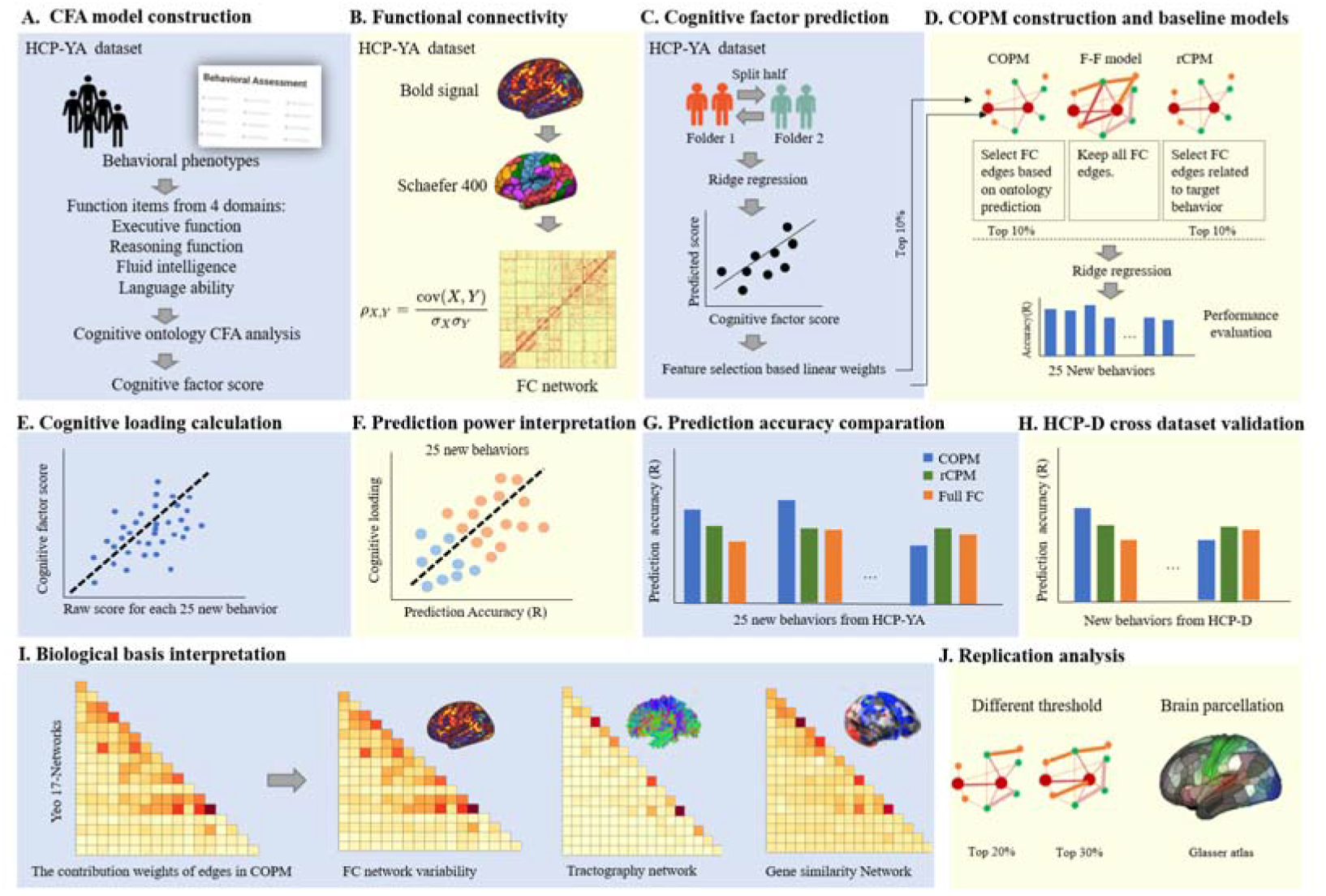
Framework of method. A – We calculated cognitive factor score at the individual level using a second-order confirmatory factor analysis (CFA), analyzing 13 behaviors across multiple domains from the HCP-YA dataset. B – We calculated functional connectivity (FC) for each participant using Schaefer 400 atlas [29]. C – We used ridge regression to predict the cognitive factor score using all FC edges. D. We then included the top 10% of edges that significantly contributed to predicting cognitive factor in the Cognitive Ontology based Prediction Model (COPM). The selected edges were then used to predict the behavior performance of 25 new behaviors from the HCP-YA dataset. To further examine whether the representation of cognitive closely related functions is distinctly tied to the above edges that play a substantial role in predicting cognitive factor score, we performed similar analyses using two alternative models. The first model, named the Full-FC (F-F) model, utilizes the entire FC matrix. The second, referred to as the Ridge-regression based Connectome Predictive Model (rCPM), used an equivalent number of edges as the COPM, but selected via multiple linear regression within the CPM framework based on their close relation to the target behavior rather than the cognitive factor. E- We calculated the correlation between each task’s raw performance and cognitive factor score across all subjects, which we refer to as ‘cognitive loading’. Tasks with higher cognitive loading indicate they measure cognitive related functions, whereas those with lower cognitive loading suggest cognitive less-related functions. Each dot presents one subject. F– We calculated the correlations between prediction accuracy calculated in D and the cognitive loading calculated in E to examine whether tasks with greater cognitive loading (i.e., those behaviors closely related with cognitive factor) have higher prediction accuracy. G – We directly compared the correlations between cognitive loading and prediction accuracy across these models. H – To examine whether the same edges identified in HCP-YA dataset also show specificity in predicting cognitive functions in a new dataset, we repeated the analysis using 8 new behaviors from the HCP-D dataset. I– To explore the biological underpinnings of the top FC edges included in the COPM, we analyzed their relationships with three key biological factors: FC variability, white matter structure, and gene expression similarity. Specifically, we extracted the contribution weights of these edges in predicting the cognitive factor score. These weights indicate each edge’s relative importance in the prediction process, with higher weights suggesting a more significant role in forecasting the cognitive factor score. We then calculate the correlations between these contribution weights and three key biological factors. J – To ensure the robustness of our findings, we replicated the main results with varying thresholds for edge selection in the COPM and employed a different brain parcellation to delineate functional areas, such as Glasser parcellation.

### The cognitive factor can be derived from a second-order CFA model and predicted by the whole brain FC

To test our hypothesis that FC edges can predict cognitive factor, we first created a cognitive factor using the behavior data and calculated FC using the resting-state fMRI data from the HCP-YA dataset. To create a cognitive factor that captures the individual variability across various cognitive domains, we employed a second-order CFA model which can more accurately capture cognitive ontology on 13 behavior tasks from the HCP-YA dataset (see Table 1). These 13 tasks span 5 domains, including: (I) 4 tasks measure executive functions related to processing speed, (II) 3 tasks measure executive functions associated with working memory, (III) 4 tasks measure fluid intelligence, (IV) 2 tasks measure relation ability, and (V) 2 tasks measure language ability. These tasks were often used to calculate the cognitive factor [2,3,10,14].

Our results showed that the five first-order factors exhibited substantial loadings in the following domains: executive function related to processing speed (β=0.47), executive function related to working memory (β=2.12), fluid intelligence (β=0.73), relational ability (β=-0.20), and language (β=0.88). Each primary factor significantly contributes to the second-order cognitive factor termed as *cognition*, encapsulating a refined representation of cognitive functions (Figure. 2A and Table. 2). The second-order CFA model can capture a second-order factor (i.e., cognitive factor) with a commendable model fit (CFI=0.95, TLI=0.93, RMSEA=0.07, 90% CI [0.06, 0.08], SRMR=0.074, χ²/df=4.16). After calculating the regression-based weights, each participant had one cognitive factor score.

**Figure 2.**
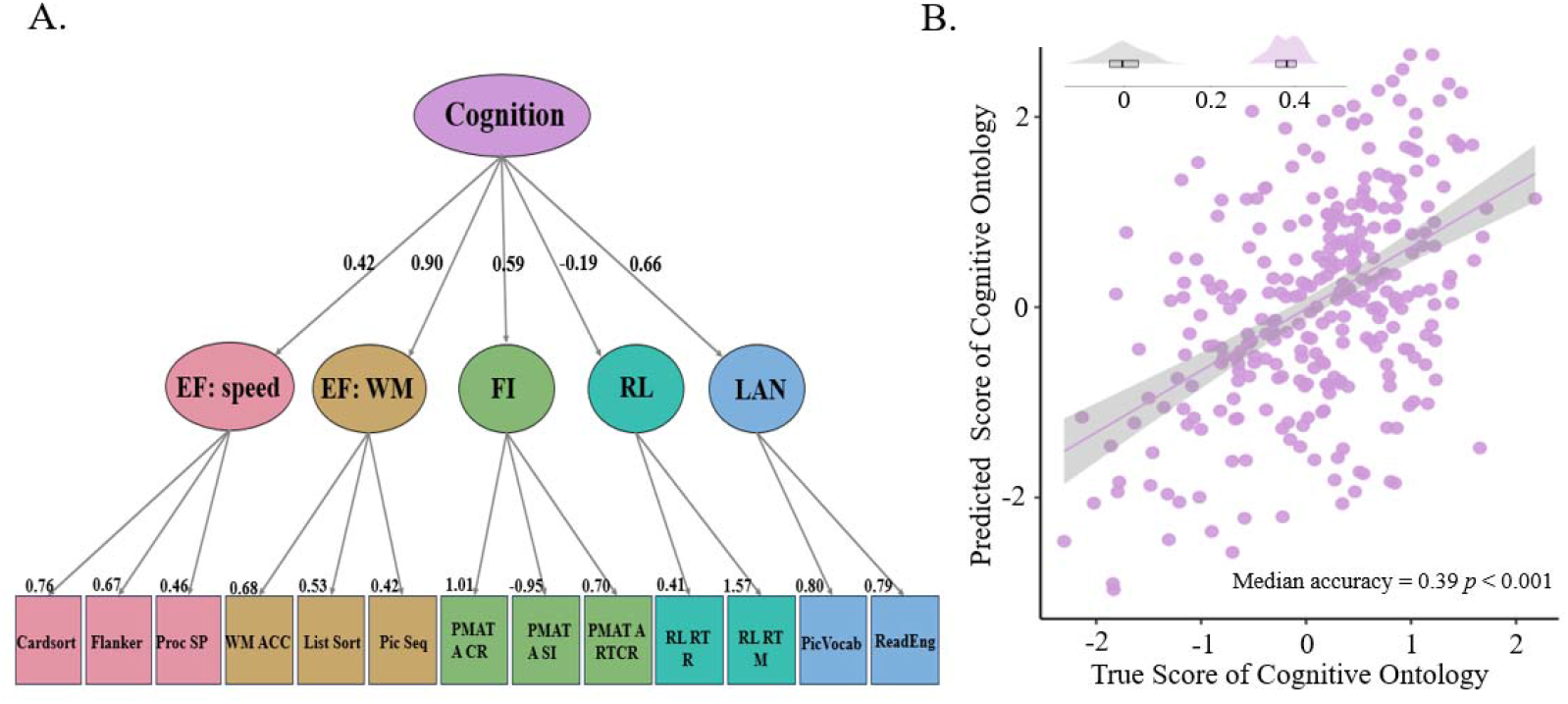
**The cognitive factor can be derived from a second-order CFA model and predicted by the whole brain FC**. A – The five first-order factors exhibited substantial loadings in the following domains: executive functions related to processing speed (EF: speed), executive functions associated with working memory (EF: WM), fluid intelligence (FI), relational ability (RL), and language (LAN). Each primary factor significantly contributes to the second-order cognitive factor, captured by the second-order CFA model. B – The cognitive factor score can be predicted by the whole brain FC with greater accuracy than chance. The scatter plot shows significant positive correlations between the actual and predicted cognitive factor scores from one iteration (the data used here was the one whose prediction accuracy most closely matched the median accuracy.). Each dot represents the data of one participant. The gray in the left corner represents the distribution of prediction accuracy from 200 permutations, while the purple represents the distribution prediction accuracy from 100 iterations of the prediction model.

Further, using the Schaefer 400 atlas, we constructed a parcel-based resting-state FC matrix for each participant and applied ridge regression to test whether the FC features can significantly predict the cognitive factor compared to the chance level (computed by permutation approach, see method ‘*(d) Prediction of the cognitive factor score using FC*’ section). We found that the whole FC matrix predicted the cognitive factor score with greater accuracy than chance (Figure 2B; p < 0.001 by nonparametric paired Wilcoxon test, with a Wilcoxon effect size of r = 0.82; the median accuracy = 0.39, interquartile range = 0.04, the median accuracy of chance level = 0.001, interquartile range of chance level = 0.067, the median MAE = 0.62, the median MAE of chance level = 0.67).

To address potential model bias, we replicated our analysis using a bifactor CFA model to derive the cognitive factor score. The bi-factor CFA model can capture a second-order factor (i.e., cognitive factor) with a perfect model fit (CFI=0.97, TLI=0.95, RMSEA=0.048, 90% CI [0.35, 0.61], SRMR=0.006, χ²/df=2.38). We observed a similar pattern that the whole FC matrix predicted the cognitive factor score with greater accuracy than chance (median accuracy = 0.41, interquartile range = 0.04, the median accuracy of chance level = 0.001, interquartile range of chance level = 0.069, the median MAE = 0.60, the median MAE of chance level = 0.67, see Figure S1 and Table S1 in Supplementary). Then we selected the top 10% edges that made significant contribution to predicting the cognitive factor for each model (i.e., second-order CFA model and bifactor CFA). We examined the consistency of the edges among the 2 models and found a strong consistency (Dice coefficient = 0.83), suggesting that FC can predict the cognitive factor, regardless of the specific CFA model used.

Additionally, to evaluate the influence of behavior selection, we adapted a G-factor model using 10 behaviors identified by Dubois et al. [10]. Among them, 7 tasks were included in the second order CFA model and 3 were new (see Table S2 for detailed information). This model also showed moderate predictive accuracy with FC (Median accuracy = 0.38, interquartile range = 0.04, median accuracy of chance level = 0.001, interquartile range of chance level = 0.071, the median MAE = 0.63, the median MAE of chance level = 0.67), and the key predictive edges largely overlapped with those from our original second-order CFA model (Dice coefficient = 0.58). These findings collectively highlight FC’s robustness in predicting cognitive factor, regardless of the CFA model or behaviors analyzed.

### The top edges that significantly contribute to the prediction of the cognitive factor represent cognitive functions more than motor and emotion functions

After identifying the top 10% of edges that contributed significantly to predicting cognitive factor score, we examined whether they could better represent cognitive functions compared to motor and emotion functions. For this purpose, we specifically selected 25 new behavioral tasks from the HCP-YA dataset to cover a spectrum of cognitive functions: some tasks, such as those measuring executive control, represented cognitive closely related functions while others, like those assessing motor skills and emotions, indicated cognitive less-related functions. This diverse range of tasks allowed us to test whether the identified edges could more accurately predict cognitive related as opposed to cognitive less-related cognitive functions. The selected tasks encompassed a broad array of cognitive and behavioral areas, including executive control, sustained attention, language, verbal episodic memory, self-regulation, spatial orientation, motor skills, emotion processing, social relationships, and psychological well-being. Detailed descriptions and comprehensive information about these tasks are provided in Table 3.

First, we calculated the correlation between each task’s raw performance and cognitive factor score across subjects (as shown in Figure 3A), which we refer to as ‘cognitive loading’. Tasks with higher cognitive loading indicate they measure cognitive related functions, whereas those with lower cognitive loading suggest they measure cognitive-less related functions. Following this, we adapted our methodology by constructing a ridge regression model to predict task performance. In this model, we deviated from our previous approach by incorporating only the top 10% of identified edges, instead of the entire FC matrix, into the COPM. After model construction, we computed the prediction accuracy for each task. Subsequently, we explored the relationship between cognitive loading and prediction accuracy by calculating their correlation. Our analysis revealed a significant positive correlation (r = 0.64, p < 0.001, Figure 3B). This finding suggests that tasks with a stronger correlation to the cognitive factor tend to have higher prediction accuracy compared to those with a weaker correlation. Consequently, these results imply that the selected edges are more representative of cognitive related functions than those less-related ones.

**Figure 3.**
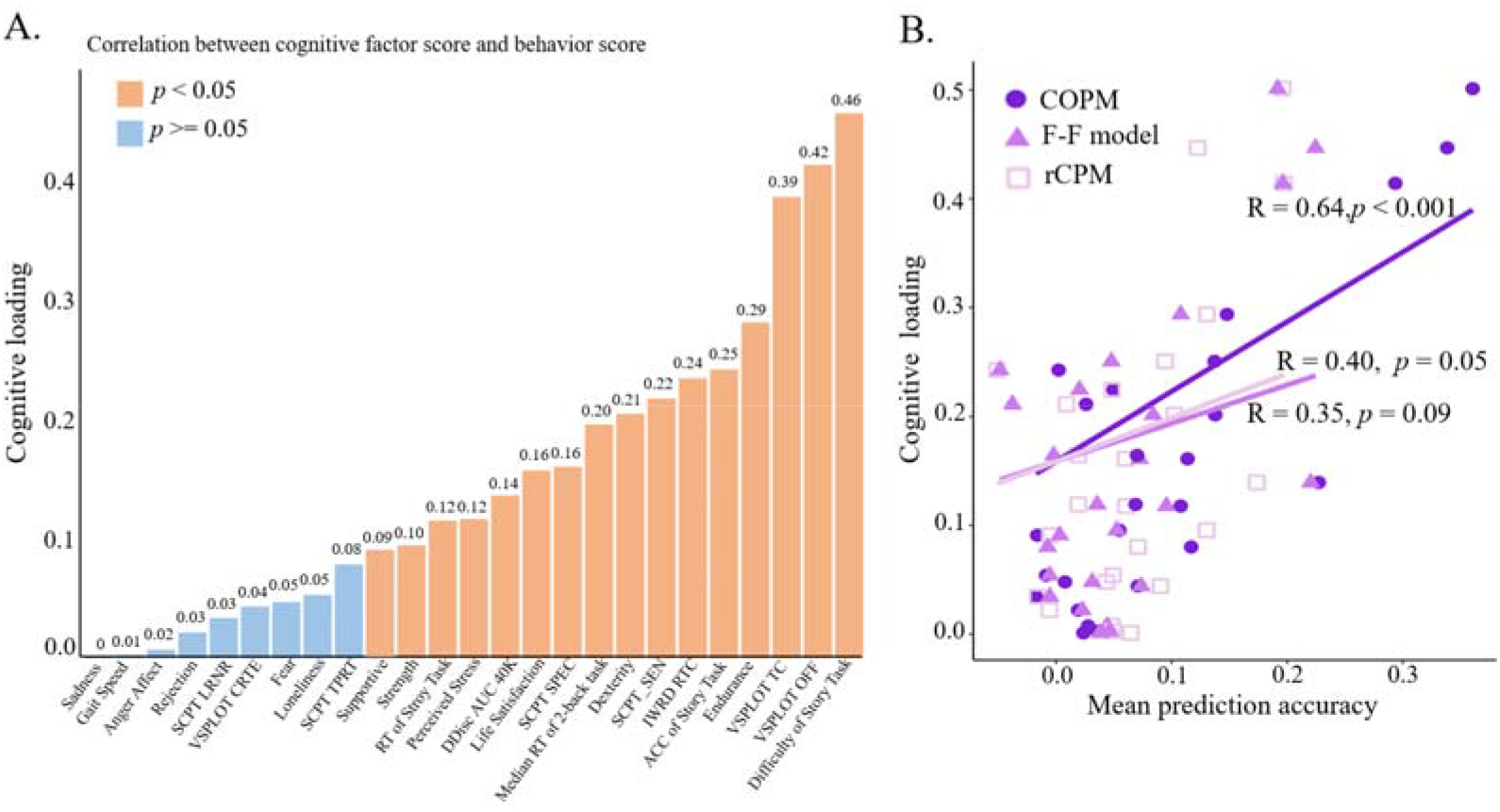
The top edges that significantly contributed to predicting cognitive factor score can better represent cognitive-related functions compared to less-related ones. A – The bar shows the correlations between each task’s raw performance and cognitive factor score across all subjects, termed ‘cognitive loading’. Tasks with higher cognitive loading indicate they measure cognitive related functions, whereas those with lower cognitive loading suggest they measure cognitive less-related functions. Orange bars represent tasks with significant correlations (p < 0.05, FDR-corrected) and Bule bars denote those with unsignificant correlations (p > 0.05, FDR-corrected). B – The scatter plot shows the correlation between cognitive loading and the prediction accuracy of each model for each task. Dark purple filled circle represents COPM, light purple filled triangle represents F-F model, and light purple empty square represents rCPM.

**Figure 4.**
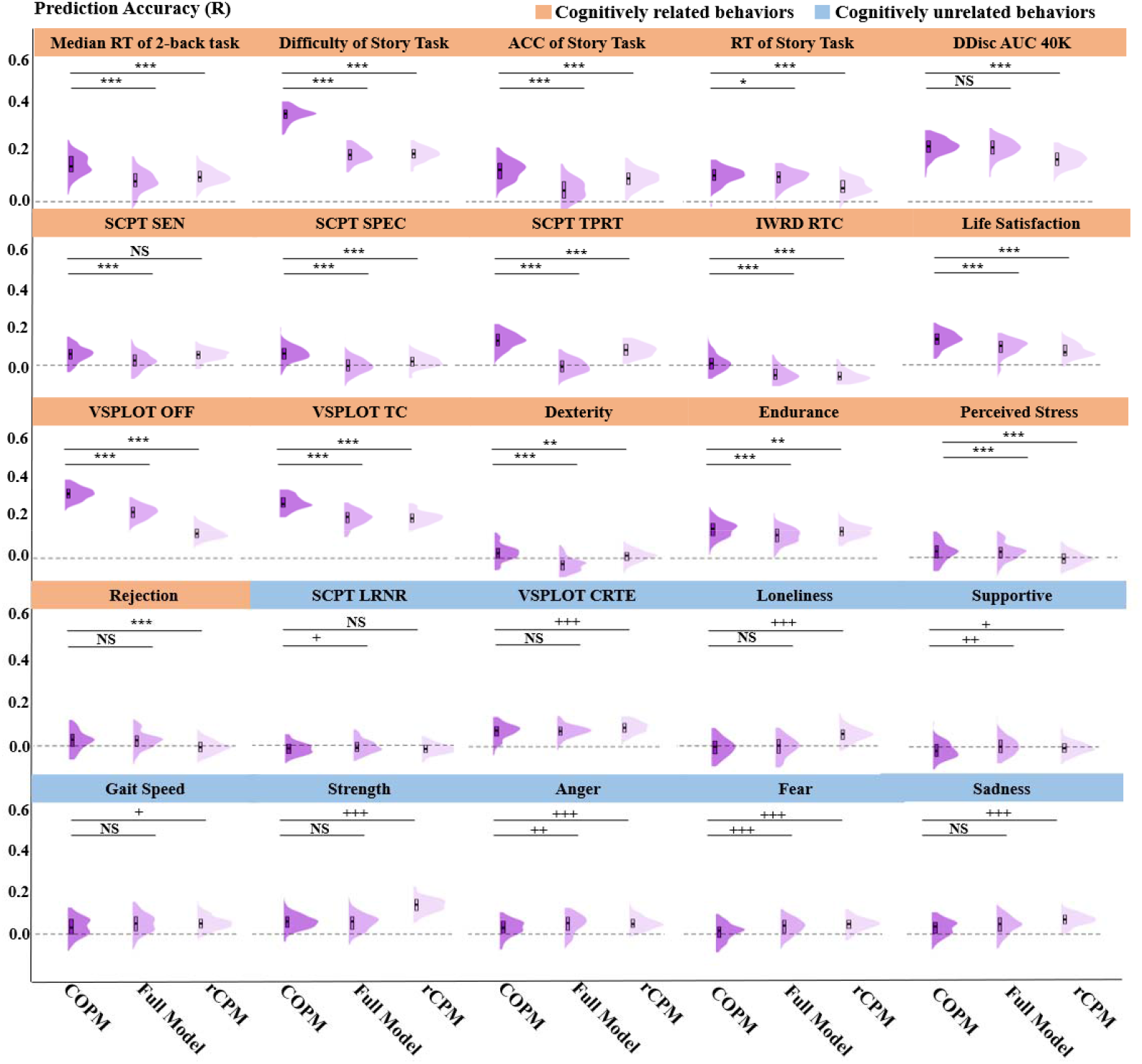
The prediction accuracy of the COPM, F-F Model and rCPM for each task in the HCP-YA dataset. Tasks whose raw performance exhibited significant correlations with the cognitive factor score are highlighted in orange. In contrast, tasks that did not show a significant correlation are highlighted in blue. The COPM demonstrates a notable ability to more accurately predict behaviors that are highly correlated with the cognitive factor score compared to the F-F and/or rCPM models. However, for behaviors with weaker correlations, the COPM exhibits a similar or even opposite prediction pattern. A * indicates that COPM has greater prediction accuracy than F-F/ rCPM. A + indicates the opposite trend. *<0.05, **<0.01, ***<0.001; +< 0.05; ++<0.01; +++<0.001; NS = not significant. All the p values are FDR-corrected. See full name of each task in Table 3.

To further examine whether the representation of cognitive function is distinctly tied to the edges that play a substantial role in predicting cognitive factor score, we performed similar analyses using 2 alternative models. The first model, named the F-F model, utilizes the entire FC matrix. The second, referred to as the rCPM, incorporates a number of edges equivalent to those in the COPM, but these edges are selected via multiple linear regression within the CPM framework. This selection is based on their close relation to the target behavior rather than the cognitive factor score (refer to the Method section for detailed information). Cognitive loading was not significantly correlated with the prediction accuracy in either the F-F model (r = 0.35, p = 0.09, Figure 3B) or the rCPM (r = 0.40, p = 0.05, Figure 3B). To further assess these findings, we directly compared the correlations between cognitive loading and prediction accuracy across these models. Cognitive loading exhibited a stronger correlation with the prediction accuracy in the COPM than in both the F-F model (z = 3.4216, *p* = 0.0006) and the rCPM (z = 3.1577, *p* = 0.0016). These results suggest that the specific edges identified as significant in predicting cognitive factor score have a unique association with cognitive functions.

To further substantiate the afore mentioned results, we categorized behaviors according to their correlation between the raw performance and the cognitive factor score. This categorization aimed to assess if the COPM could predict highly correlated behaviors more accurately than the other 2 models and if it demonstrated a similar or opposite pattern for behaviors with weaker correlations. Our findings indicated that 16 out of 25 behaviors showed significant correlations with the cognitive factor score (p < 0.05, FDR-corrected), whereas 9 did not. The COPM predicted the performance of the 16 significantly correlated behaviors more accurately than the 9 with weaker correlations (p < 0.001, Wilcoxon effect size r = 0.868). Within the group of behaviors demonstrating significant correlations with cognitive factor, the COPM outperformed both the F-F Model and the rCPM in predicting the performance of 13 behaviors (p<0.05, FDR corrected, see details in table 4). These behaviors included tasks measuring executive function and language abilities, such as the reaction time (RT) of 2-back working memory and the accuracy of the story comprehension task. In contrast, for 2 behaviors in this subset, the COPM yielded greater prediction accuracies than either the F-F Model or rCPM, including tasks like the RT of the story comprehension task. Besides, the prediction accuracy of strength task of COPM lower than rCPM. However, for the subset of behaviors that did not show significant correlations, the predictive accuracy of the COPM was exceeded by either the F-F Model or the rCPM except the median RT for true response in Short Penn Continuous Performance Test, which evaluates cognitive speed in attention task (p > 0.05, FDR-corrected).

### The same edges identified in the HCP-YA dataset also show specificity in predicting cognitive related functions in the HCP-D dataset

We then validated our findings using the HCP-D dataset. We selected 6 tasks designed to assess cognitive functions, including executive functions and language abilities, along with 2 tasks aimed at evaluating motor and emotion function (see Table 1 and Table 2 for detailed descriptions). We then compared the prediction accuracies across the 2 types of behaviors and the three models. We found that COPM predicted the performance of cognitive related function tasks with significantly greater accuracy than those for cognitive less-related functions (p < 0.001, Wilcoxon effect size r = 0.868). Within the cognitive related function tasks, the COPM outperformed both the F-F Model and the rCPM in 5 out of 6 tasks (p < 0.05, FDR corrected; see Figure 5 and Table 5). However, for 2 tasks assessing cognitive less-related functions, the prediction accuracies of the COPM, the F-F Model, and the rCPM were comparable (p > 0.05, FDR-corrected; Figure 5 and Table 5).

**Tabel 5.** The prediction accuracy of all 8 behaviors was assessed using a cross-dataset prediction paradigm for the three models.

| Domains | Behavior | Full FC (SD) | rCPM (SD) | COPM (SD) |
| --- | --- | --- | --- | --- |
| Executive function | DCCS | 0.036 (0.052) | 0.037 (0.052) | 0.046 (0.050) |
| Executive function | Flanker | -0.045 (0.058) | -0.056 (0.057) | 0.005 (0.057) |
| Executive function | Picture sequence memory | 0.002(0.054) | 0.005 (0.052) | 0.024 (0.045) |
| Executive function | List sorting working memory | 0.071(0.051) | 0.065 (0.053) | 0.111 (0.050) |
| Language | Picture vocabulary | 0.236 (0.040) | 0.220 (0.041) | 0.267 (0.041) |
| Language | Oral reading recognition | 0.169 (0.051) | 0.151 (0.050) | 0.186 (0.050) |
| Emotion | Fear | -0.127 (0.050) | -0.118 (0.050) | -0.123 (0.050) |
| Motor | Strength | 0.025 (0.058) | 0.029 (0.056) | 0.013 (0.054) |

**Figure 5.**
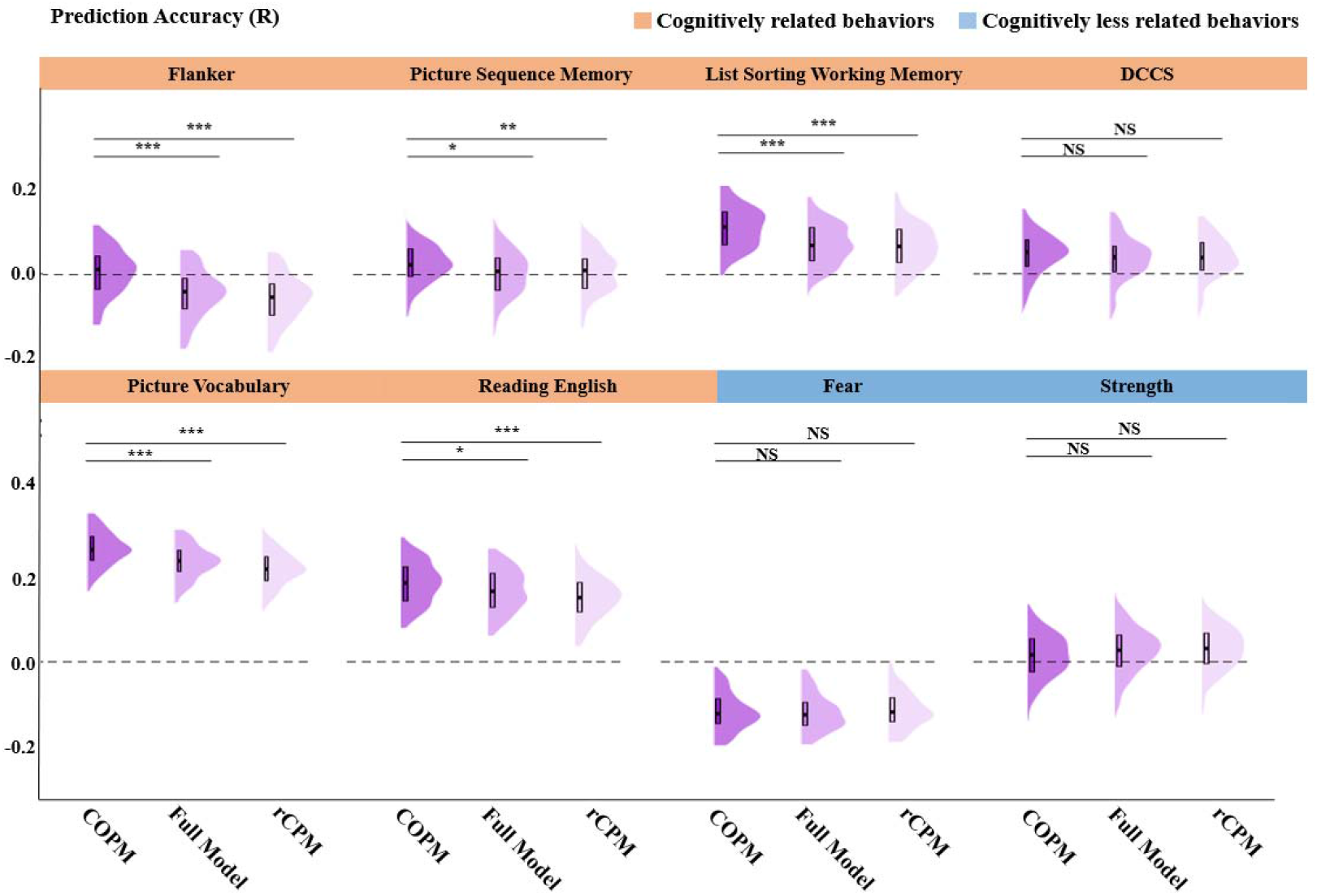
The prediction accuracy of the COPM, F-F Model and rCPM for each task in the HCPD dataset. Tasks whose raw performance exhibited significant correlations with the cognitive factor score in HCP-YA are highlighted in orange. In contrast, tasks that did not show a significant correlation are highlighted in blue. The COPM demonstrates a notable ability to more accurately predict behaviors that are highly correlated with the cognitive factor score compared to the F-F and/or rCPM models. However, for behaviors with weaker correlations, the COPM exhibits a similar or even opposite prediction pattern. A * indicates that COPM has greater prediction accuracy than F-F/ rCPM. *<0.05, **<0.01, ***<0.001; +< 0.05; NS = not significant. All the p values are FDR-corrected.

### Edges with higher predictive contributions to ontology scores show greater FC variability, stronger SC, and more similar gene expression profiles

Subsequently, we explored the biological underpinnings of the top 10% FC edges included in the COPM. Specifically, we extracted the contribution weights of these edges in predicting the cognitive factor. These weights indicate each edge’s relative importance in the prediction process, with higher weights suggesting a more significant role in forecasting the cognitive factor score. We then investigated the relationship between these contribution weights and three key biological factors: individual variability in FC strength, structural connectivity (SC) strength, and gene expression similarity profiles. Previous research found the SC-FC coupling across distinct brain regions supports the cognitive process, and SC–FC coupling is heritable[43], thus, we hypothesized the edges that made greater contribution when predicting the cognitive factor would show greater individual variability in FC strength, stronger SC, and more similar gene expression profiles.

To assess the individual variability of each FC edge, we calculated the ratio of inter-individual variability to intra-individual variability for each edge. This method helps offset the influence of FC magnitude on variability assessments, as connectivity strength tends to affect both inter- and intra-individual variability similarly (i.e., edges with stronger FC typically exhibit greater variability on both counts). As a result, we generated a symmetrical 400x400 matrix, with each edge representing inter-individual variability across a cohort of 601 subjects from HCP-YA dataset. In parallel with FC analysis, we also measured SC strength using diffusion MRI data from the HCP-YA dataset to create a white matter structural connectivity matrix. We constructed a 400x400 SC matrix for each participant and then computed the average across all participants. Furthermore, we measured the gene expression similarity profiles using the Allen Human Brain Atlas which is a transcriptional atlas [44]. This atlas includes gene expression levels measured with DNA microarray probes and sampled from hundreds of neuroanatomical structures in the left hemisphere across 4 normal postmortem human brains and bilateral hemisphere across 2 additional human brains (Sunkin et al., 2013; Burt et al., 2018). From these data, we calculated group-averaged gene expression profiles across 400 cortical areas using Schaefer parcellation [29].

In our analysis, we focused solely on the edges included in the COPM. For each measurement, we averaged the estimates both within and between networks to create a network-by-network matrix, as outlined in the methodologies by Schaefer et al. (2018). We then computed Spearman correlations to explore the relationship between the contribution weights of these edges and their corresponding metrics in the three matrices: FC variability, SC strength, and gene expression profiles. We found that the contribution weights were significantly correlated with the FC variability (r = 0.96, p < 0.001, FDR corrected), SC strength (r = 0.59, p < 0.001, FDR corrected), and gene expression profiles (r = 0.54, p = 0.002, FDR corrected). These results indicate that the edges that made greater contribution when predicting the cognitive factor score also showed greater variability in FC, stronger SC, and more similar gene expression profiles. As shown in Figure 6, edges making greater contributions to predicting cognitive factor are mainly located within the association networks, particularly within the intra-and inter-connections of the Default mode network (DMN) and Frontal-parietal control network (FPCN).

**Figure 6.**
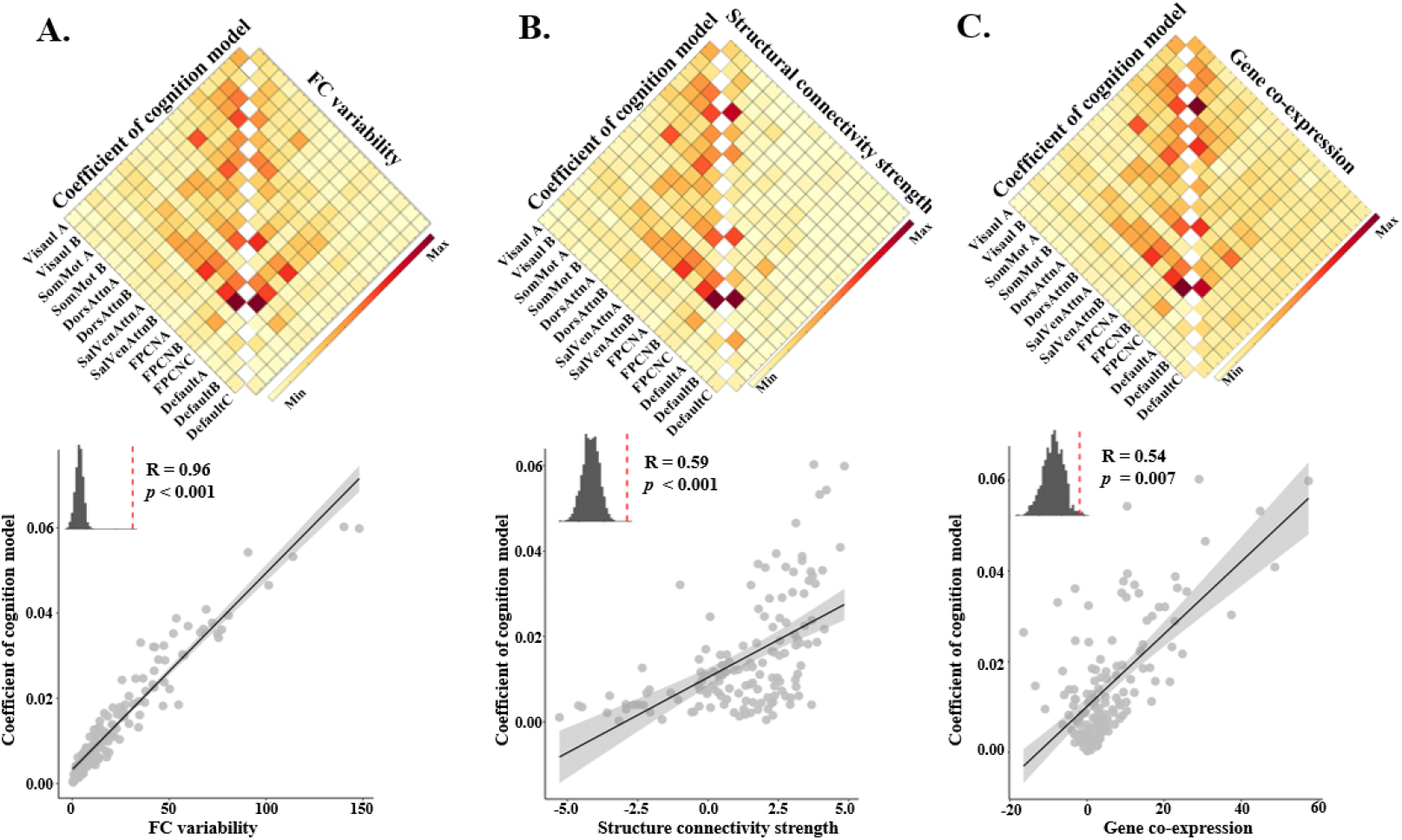
Edges that made greater contribution when predicting the cognitive factor score showed greater variability in FC, stronger SC, and more similar gene expression profiles. A, B, and C show the significant Spearman correlations between the contribution weights of the edges in predicting the cognitive factor score and their corresponding metrics in the three matrices: FC variability, structural connectivity strength, and gene expression profiles (p < 0.05, FDR-corrected).

### Our findings are robust when using varied thresholds for edge selection and Glasser atlas

To evaluate the robustness of our results, we experimented with varying thresholds for edge selection within the COPM, alongside using different brain parcellations to delineate functional areas. While our primary analysis focused on the top 10% of edges, we expanded this to include thresholds of 20%, 30%. For each threshold, we predicted task-specific behavioral performance and analyzed the correlation between cognitive loading and prediction accuracy. We consistently observed significant correlations at these extended thresholds (20%: r = 0.57, p = 0.002; 30%: r =0.51, p = 0.005). These results reinforce our initial findings, indicating that tasks with a high correlation to cognitive factor consistently demonstrate enhanced prediction accuracy. This trend persists across various thresholds, suggesting a reliable and accurate representation of cognitive related functions.

Furthermore, we conducted parallel analyses using the rCPM, which selects an equivalent number of edges as the COPM but specifically targets edges associated with each behavior rather than cognitive factor score using multiple linear regression within the CPM framework. Cognitive loading also showed significant correlations with prediction accuracy in the rCPM across the varied thresholds (20%: r = 0.46, p = 0.02, 30%: r = 0.46, p = 0.02). However, when directly comparing the 2 models, the correlation between cognitive loading and prediction accuracy was stronger in the COPM than in the rCPM at the 20% and 30% thresholds (20%: z = 2.50, p = 0.01, 30%: z = 2.27, p = 0.02). This pattern suggests that the edges selected by the COPM, particularly at lower thresholds, are distinctively associated with the representation of cognitive related functions. Overall, these findings highlight the COPM’s effectiveness in utilizing a reduced set of FC edges for more precise predictions related to cognitive factor.

After confirming that our results were not influenced by the threshold used, we replicated our analysis using Glasser atlas (Glasser et al., 2016), applying a 10% threshold. Given the comparable predictive efficiency of the rCPM and the F-F model, and due to computational and time constraints, we limited our analysis to the F-F and COPM when using the Glasser 360 atlas. The FC network built with the Glasser 360 atlas showed moderate predictive accuracy for the cognitive factor factor (R median = 0.29, interquartile range = 0.05).

We observed a significant correlation between cognitive loading and prediction accuracy in the COPM (r = 0.63, p = 0.002). This finding reinforces our earlier observation: tasks with a closer relationship to the cognitive factor tend to have higher prediction accuracy than those with weaker correlations. Such a correlation was not evident in the F-F model (r =0.40, p = 0.10). The stronger correlation in the COPM compared to the F-F model (z =2.8814, p = 0.004) further substantiates the notion that specific predictive edges in the COPM are uniquely linked to cognitive related functions, a linkage that remains consistent across different brain atlases. Owing to the limitations in computational resources and time, we did not extend this replication to other thresholds or atlases.

## Discussion

This study identified a set of core FC connectomes, predominantly located within the association cortex of the human brain, that represent cognition. Leveraging a novel machine learning framework, the COPM, these core FC connectomes not only predict the cognitive factor score with high accuracy but also efficiently forecast a spectrum of cognitive functions, affirming their applicability across datasets. The core FC connectome edges are characterized by substantial inter-individual variability in FC strength, strong SC, and comparable gene expression patterns. Our findings remained robust across various edge selection thresholds and the application of the Glasser atlas.

Our research not only replicated previous findings on the predictive capacity of FC edges for the cognitive factor [10–12,20] but also expanded them by pinpointing a specific set of FC edges within association cortex networks like the DMN, FPCN, and attention network. This discovery aligns with existing findings on the necessity of coordinated activities across distributed brain regions, primarily within the association cortex, for numerous cognitive functions [6,9,45,46]. Moreover, our findings support the massive redeployment hypothesis [47] suggesting a universal brain region framework underpinning diverse cognitive functions.

We also extended previous findings by exploring the biological foundations of these key edges, encompassing aspects like FC variability, white matter integrity, and gene expression similarity. We observed that these edges exhibit not only a high degree of variability in FC but also strong SC and closely matched gene expression profiles. This increased FC variability suggests a potential for enhanced neural adaptability, providing the neural flexibility needed for various cognitive tasks [40,48]. Such findings highlight the brain’s dynamic nature and its capacity for plasticity – the ability to modify connections based on experiences, information, or acquired skills. In line with this, the strength of FC within the executive control network have been shown to be positively correlated with performance in sustained cognitive performance [19,49]. This correlation underscores the importance of these connections in learning and cognitive adaptation. Additionally, the association cortex, known for integrating information across diverse brain areas, plays a vital role in complex cognitive functions like decision-making and problem-solving. This integrative capability may underlie the remarkable cognitive flexibility humans demonstrate, allowing us to respond innovatively and adaptively to a constantly changing environment [46]. The observation that certain FC edges exhibited stronger SC and similar gene expression patterns hints at a genetic foundation for these neural connections. This suggests that genetic factors may influence the formation and maintenance of brain networks, affecting how regions connect and interact, and how they are shaped by experiences [50,51].

A key novelty of our study is the COPM, a cognitive ontology-based machine learning framework. Distinct from traditional approaches that utilize all edges in predicting the cognitive factor, the COPM identifies critical edges that are predictive of the cognitive factor. These selected edges can predict a wide range of cognitive functions, such as executive function, language ability, self-regulation, sustained attention, verbal episodic memory, and spatial orientation, but not motor and emotion functions. This approach holds promise for uncovering robust biomarkers specific to cognitive tasks or impairments, offering potential for refined diagnostic and intervention strategies in clinical contexts [9,33,52,53].

Despite its contributions, our study has methodological limitations. Firstly, our analysis of FC edges’ importance relied on linear model’s coefficient weights, a technique paralleled in prior research [17,54–56]. However, this method may not fully capture the significance of FC edges. Future investigation could explore alternative techniques, such as feature dropout [10]or edge permutation importance [20], to more accurately evaluate the edges’ impact on model performance. Secondly, while we have established correlations among core FC networks, gene expression patterns, and cognitive factors, a direct exploration of causality between structural networks, gene expression and cognitive processes could yield deeper insights into the brain’s cognitive architecture. For example, machine learning algorithms could be applied to multimodal brain imaging and genetics data to predict cognition scores. Structural equation modeling may also clarify the directionality of these relationships.

## Conclusion

This study identifies and validates a set of functional connectomes within the association cortex that precisely represents cognition, as evidenced by the COPM. These core connectomes demonstrate superior predictive accuracy for various cognitive functions, such as working memory, reading comprehension, and sustained attention, and characterized by significant individual variability in FC strength, interconnected via white matter tracts, and gene expression similarities.

## Acknowledgments

The authors thank Hu Chuan Peng for helping with advisement. Funding: STI 2030—Major Projects 2021ZD0201500 (YD); National Natural Science Foundation of China 31822024 (YD) and 32300881 (XYW); Strategic Priority Research Program of Chinese Academy of Sciences XDB32010300 (YD); Scientific Foundation of Institute of Psychology, Chinese Academy of Sciences E2CX3625CX (Y.D.) and E1CX4725CX (XYW). Data were provided by the Human Connectome Project, WU-Minn Consortium (Principal Investigators: David Van Essen and Kamil Ugurbil; 1U54MH091657) funded by the 16 NIH Institutes and Centers that support the NIH Blueprint for Neuroscience Research; and by the McDonnell Center for Systems Neuroscience at Washington University. The HCP-Development 2.0 Release data used in this report came from DOI: 10.15154/1520708. Research reported in this publication was supported by the National Institute of Mental Health of the National Institutes of Health under Award Number U01MH109589 and by funds provided by the McDonnell Center for Systems Neuroscience at Washington University in St. Louis.

## Notes

https://osf.io/zu3nt/?view_only=85483542699c41db8b48a155dad5cb7a

### Competing Interest Statement

The authors have declared no competing interest.

### Summary of Updates

Change fortmat of author's name to correct format.

