## Supplemental Table and Figures for "Unveiling the Core Functional Networks of Cognition: An Ontology-Guided Machine Learning Approach"

**Supplementary Materials**

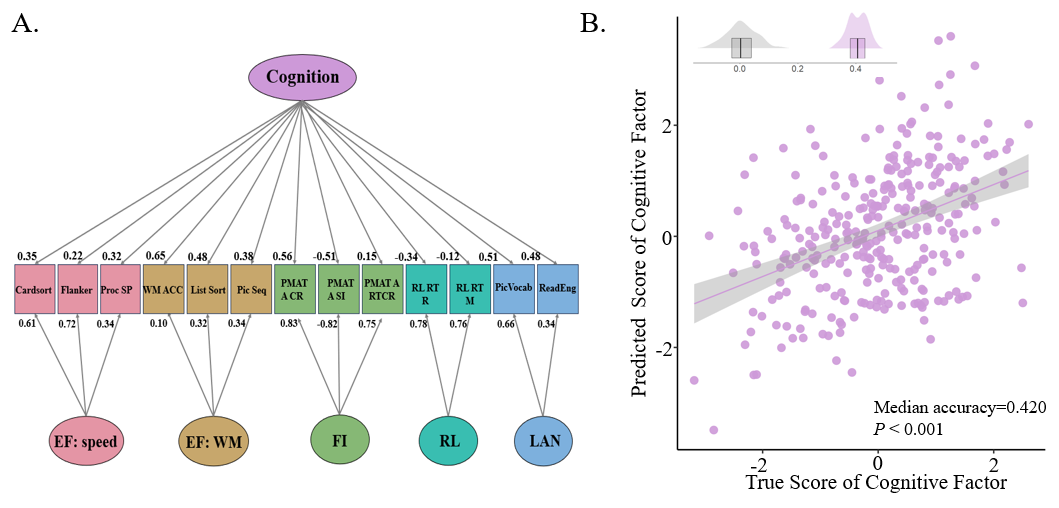

Figure S1 The bifactor CFA model and the prediction accuracy for the cognitive ontology score from bifactor model. A – The five group factors exhibited substantial loadings in the following domains: executive functions related to processing speed (EF: speed), executive functions associated with working memory (EF: WM), fluid intelligence (FI), relational ability (RL), and language (LAN). Each behavioral task significantly contributes to the general factor: cognitive factor, captured by the second-order CFA model. B – The cognitive factor score can be predicted by the whole brain FC with greater accuracy than chance. The scatter plot shows significant positive correlations between the actual and predicted cognitive factor scores from one iteration (the data used here was the one whose prediction accuracy most closely matched the median accuracy.). Each dot represents the data of one participant. The gray in the left corner represents the distribution of prediction accuracy from 200 permutations, while the purple represents the distribution prediction accuracy from 100 iterations of the prediction model.

Table S1 Fitting results of bi-factor CFA model

| Latent Variables | Estimate (S.E.) | z | p | LCI | UCI |
| --- | --- | --- | --- | --- | --- |
| EF: Speed =~ Card Sorting | 0.61 (0.06) | 9.67 | <.001 *** | 0.48 | 0.72 |
| EF: Speed =~ Flanker | 0.72 (0.07) | 10.23 | <.001 *** | 0.57 | 0.84 |
| EF: Speed =~ Process Speed | 0.34 (0.05) | 6.95 | <.001 *** | 0.24 | 0.43 |
| EF: WM =~ WM Task 2bk Acc | 0.11 (0.09) | 1.11 | 0.27 | -0.08 | 0.27 |
| EF: WM =~ List Sorting | 0.32 (0.22) | 1.43 | 0.15 | -0.12 | 0.75 |
| EF: WM =~ Picture Sequence | 0.34 (0.24) | 1.42 | 0.16 | -0.13 | 0.81 |
| FI =~ PMAT24 A CR | 0.83 (0.03) | 26.38 | <.001 *** | 0.76 | 0.88 |
| FI =~ PMAT24 A SI | -0.82 (0.03) | -25.38 | <.001 *** | -0.87 | -0.75 |
| FI =~ PMAT24 A RTCR | 0.75 (0.04) | 20.41 | <.001 *** | 0.67 | 0.82 |
| Relation =~ Match Median RT | 0.78 (46.65) | 0.02 | 0.99 | -90.65 | 92.21 |
| Relation =~ Rel Median RT | 0.76 (45.62) | 0.02 | 0.99 | -88.66 | 90.18 |
| LAN =~ Pic Vocabulary | 0.65 (44.91) | 0.01 | 0.99 | -87.37 | 88.66 |
| LAN =~ Read English | 0.34 (40.99) | 0.01 | 0.99 | -79.76 | 80.93 |
| Cognition =~ Card Sorting | 0.35 (0.05) | 7.44 | <.001 *** | 0.26 | 0.44 |
| Cognition =~ Flanker | 0.22 (0.05) | 4.63 | <.001 *** | 0.13 | 0.31 |
| Cognition =~ Process Speed | 0.32 (0.05) | 6.74 | <.001 *** | 0.23 | 0.41 |
| Cognition =~ WM Task 2bk Acc | 0.64 (0.05) | 13.52 | <.001 *** | 0.54 | 0.73 |
| Cognition =~ List Sorting | 0.48 (0.05) | 9.75 | <.001 *** | 0.38 | 0.57 |
| Cognition =~ Picture Sequence | 0.37 (0.05) | 7.37 | <.001 *** | 0.27 | 0.47 |
| Cognition =~ PMAT24 A CR | 0.55 (0.05) | 11.53 | <.001 *** | 0.46 | 0.64 |
| Cognition =~ PMAT24 A SI | -0.50 (0.05) | -10.41 | <.001 *** | -0.59 | -0.41 |
| Cognition =~ PMAT24 A RTCR | 0.15 (0.05) | 2.91 | <.001 *** | 0.05 | 0.25 |
| Cognition =~ Match Median RT | -0.34 (0.05) | -7.14 | <.001 *** | -0.44 | -0.25 |
| Cognition =~ Rel Median RT | -0.12 (0.05) | -2.41 | 0.016 * | -0.21 | -0.02 |
| Cognition =~ Pic Vocabulary | 0.50 (0.05) | 10.81 | <.001 *** | 0.41 | 0.59 |
| Cognition =~ Read English | 0.48 (0.05) | 10.24 | <.001 *** | 0.38 | 0.57 |

Note: A * indicates p<0.05, two * indicates p<0.01 and three indicates p<0.001, LCI means lower bound of confidence interval and UCI means upper bound of confidence interval

Table S2 10 behaviors used in g-factor model

| **Test Name** | **Test Content** |
| --- | --- |
| NIH toolbox:  DCCS | Dimensional Change Card Sort is a measure of cognitive flexibility. Score is based on a combination of accuracy and reaction time, and the test takes approximately 4 minutes to administer |
| NIH toolbox:  Flanker | The Flanker task measures both a participant's attention and inhibitory control. Scoring is based on a combination of accuracy and reaction time, and the test takes approximately 3 minutes to administer |
| NIH toolbox:  Process Speed | Pattern Completion Processing Speed measures speed of processing. Asking participants to discern whether 2 side-by-side pictures are the same or not. Participants’ raw score is the number of items correct in a 90-second period. |
| NIH toolbox:  List Sorting | List Sorting assesses working memory and requires the participant to sequence different visually- and orally-presented stimuli. |
| NIH toolbox:  Picture Sequence | Picture Sequence Memory involves recalling increasingly lengthy series of illustrated objects and activities that are presented in a particular order on the computer screen. |
| PMAT24 A CR | Penn Progressive Matrices: Number of Correct Responses |
| VSPLOT_TC | Total completion in Penn Line Orientation Test. Assesses completion rate in visual-spatial task |
| IWRD_TOT | Penn Word Memory Test: Total Number of Correct Responses. |
| NIH toolbox:  Picture Vocabulary | The respondent is presented with an audio recording of a word and 4 photographic images on the computer screen and is asked to select the picture that most closely matches the meaning of the word. |
| NIH toolbox:  Reading Recognition | The participant is asked to read and pronounce letters and words as accurately as possible. |

Table S3 The effective size (Wilcoxon effect size r) and *p* value for the comparative results between COPM and F-F model, rCPM for behaviors from HCP-A dataset

| Domains | Behavior | Full FC (*p*) | rCPM (*p*) |
| --- | --- | --- | --- |
| Executive function | WM Task 2bk Median RT | 0.695 (<0.001) | 0.557 (<0.001) |
| Emotion | Anger Affect | -0.279 (0.008) | -0.385 (<0.001) |
| Emotion | Sadness | -0.190 (0.075) | -0.667 (<0.001) |
| Emotion | Fear Affect | -0.364 (<0.001) | -0.859 (<0.001) |
| Language | Language Task Story Avg Difficulty | 0.868 (<0.001) | 0.868 (<0.001) |
| Language | Language Task Story Acc | 0.833 (<0.001) | 0.632 (<0.001) |
| Language | Language Task Story Median RT | 0.240 (0.023) | 0.711 (<0.001) |
| Psychological Well-being | Life Satisfy | 0.631 (<0.001) | 0.737 (<0.001) |
| Motor | Endurance | 0.519 (<0.001) | 0.286 (0.005) |
| Motor | Strength | 0.036 (0.752) | 0.833 (<0.001) |
| Motor | Gait Speed Comp | 0.148 (0.175) | 0.203 (0.047) |
| Motor | Dexterity | 0.770 (<0.001) | 0.277 (<0.007) |
| Sustained Attention | SCPT TPRT | 0.866 (<0.001) | 0.855 (<0.001) |
| Sustained Attention | SCPT SPEC | 0.809 (<0.001) | 0.722 (<0.001) |
| Sustained Attention | SCPT SEN | 0.488 (<0.001) | 0.022 (0.827) |
| Sustained Attention | SCPT LRNR | -0.264 (0.012) | -0.060 (0.571) |
| Spatial Orientation | VSPLOT OFF | 0.866 (<0.001) | 0.868 (<0.001) |
| Spatial Orientation | VSPLOT TC | 0.857 (<0.001) | 0.862 (<0.001) |
| Spatial Orientation | VSPLOT CRTE | -0.010 (0.919) | -0.428 (<0.001) |
| Self-Regulation | DDisc AUC 40K | 0.137 (0.206) | 0.728 (<0.001) |
| Social Relationship | Perc Reject | 0.097 (0.379) | 0.428 (<0.001) |
| Social Relationship | Loneliness | 0.044 (0.719) | 0.763 (<0.001) |
| Social Relationship | Instru Supp | -0.321 (0.002) | -0.232 (0.023) |
| Social Relationship | PercStress | 0.552 (<0.001) | 0.711 (<0.001) |
| Verbal Episodic Memory | IWRD RTC | 0.661 (<0.001) | 0.740 (<0.001) |

Note: positive effective size indicates that the prediction accuracy of COPM larger than F-F model or rCPM, and negative effective size indicates that the prediction accuracy of F-F model or rCPM larger than COPM. All *p* values are corrected by FDR.

TableS4 The effective size (Wilcoxon effect size r) and *p* value for the comparative results between COPM and F-F model, rCPM for behaviors from HCP-D dataset

| Domains | Behavior | Full FC (SD) | rCPM (SD) |
| --- | --- | --- | --- |
| Executive function | DCCS | 0.135 (0.204) | 0.129 (0.226) |
| Executive function | Flanker | 0.537 (<0.001) | 0.628 (<0.001) |
| Executive function | Picture sequence memory | 0.290 (0.008) | 0.264 (0.013) |
| Executive function | List sorting working memory | 0.527 (<0.001) | 0.548 (<0.001) |
| Language | Picture vocabulary | 0.500 (<0.001) | 0.638 (<0.001) |
| Language | Oral reading recognition | 0.247 (0.022) | 0.406 (<0.001) |
| Emotion | Fear | 0.041 (0.684) | -0.090 (0.370) |
| Motor | Strength | -0.204 (0.056) | -0.199 (0.063) |

Note: positive effective size indicates that the prediction accuracy of COPM larger than F-F model or rCPM, and negative effective size indicates that the prediction accuracy of F-F model or rCPM larger than COPM. All *p* values are corrected by FDR.
